# Tepsin binds LC3B to promote ATG9A export and delivery at the cell periphery

**DOI:** 10.1101/2023.07.18.549521

**Authors:** Natalie S. Wallace, John E. Gadbery, Cameron I. Cohen, Amy K. Kendall, Lauren P. Jackson

## Abstract

Tepsin is an established accessory protein found in Adaptor Protein 4 (AP-4) coated vesicles, but the biological role of tepsin remains unknown. AP-4 vesicles originate at the *trans*-Golgi network (TGN) and target the delivery of ATG9A, a scramblase required for autophagosome biogenesis, to the cell periphery. Using *in silico* methods, we identified a putative <u>L</u>C3-Interacting <u>R</u>egion (LIR) motif in tepsin. Biochemical experiments using purified recombinant proteins indicate tepsin directly binds LC3B, but not other members, of the mammalian ATG8 family. Calorimetry and structural modeling data indicate this interaction occurs with micromolar affinity using the established LC3B LIR docking site. Loss of tepsin in cultured cells dysregulates ATG9A export from the TGN as well as ATG9A distribution at the cell periphery. Tepsin depletion in a mRFP-GFP-LC3B HeLa reporter cell line using siRNA knockdown increases autophagosome volume and number, but does not appear to affect flux through the autophagic pathway. Re-introduction of wild-type tepsin partially rescues ATG9A cargo trafficking defects. In contrast, re-introducing tepsin with a mutated LIR motif or missing N-terminus does not fully rescue altered ATG9A subcellular distribution. Together, these data suggest roles for tepsin in cargo export from the TGN; delivery of ATG9A-positive vesicles at the cell periphery; and in overall maintenance of autophagosome structure.

## Introduction

Membrane trafficking pathways are fundamental for diverse cellular and physiological functions. The assembly of cytosolic coat protein complexes carefully regulates vesicle and tubule transport between compartments in the secretory and post-Golgi trafficking networks (Dacks and Robinson, 2017). Mammalian AP (<u>A</u>ssembly <u>P</u>olypeptide) complexes 1-5 are a family of structurally homologous heterotetramers that drive vesicle coat formation at various organelle membranes (Sanger et al., 2019). The structures, mechanisms, and functions of coat formation are well understood for clathrin-associated complexes AP-1 and AP-2 (Robinson, 2015). In contrast, non-clathrin-associated AP coats are less well understood, with some exhibiting unique compositions (AP-5) and different assembly mechanisms (Hirst et al., 2021; Schoppe et al., 2020, 2021). The AP-4 complex (ε/β4/μ4/−4 subunits) is recruited to the *trans*-Golgi network (TGN) by Arf1 (Boehm et al., 2001). AP-4 does not interact with clathrin (Dell’Angelica et al., 1999; Hirst et al., 1999), and no scaffold proteins have been identified in AP-4 coated vesicles. AP-4 is ubiquitously expressed but appears particularly important in neurons and neuronal tissues. AP-4 loss in humans results in a complex neurological disorder termed AP-4-deficiency syndrome (Ebrahimi-Fakhari et al., 1993; Jameel et al., 2014; Abdollahpour et al., 2014; Abou Jamra et al., 2011; Bauer et al., 2012; Hardies et al., 2015; Tessa et al., 2016).

Biochemical approaches including proteomics have identified AP-4 coat accessory proteins: tepsin (Mattera et al., 2015; Frazier et al., 2016; Borner et al., 2012); RUSC1 and RUSC2 (Davies et al., 2018); and Hook1 and Hook2 (Mattera et al., 2020). These studies also identified several AP-4 transmembrane protein cargoes: ATG9A (Mattera et al., 2017; Davies et al., 2018; Ivankovic et al., 2020), SERINC1 and SERINC3 (Davies et al., 2018), and diacylglycerol lipase β, or DAGLB (Davies et al., 2022). ATG9A trafficking by AP-4 has been identified as a diagnostic marker of AP-4- deficiency syndrome to aid development of therapeutics (Behne et al., 2020). This discovery highlights the importance of revealing the molecular mechanisms of AP-4 biology. Loss of AP-4 in many cell types, including patient-derived cells (Davies et al., 2018; Behne et al., 2020), results in retention of ATG9A in the TGN (Mattera et al., 2017; Ivankovic et al., 2020). ATG9A is a lipid scramblase (Maeda et al., 2020; Matoba et al., 2020; Guardia et al., 2020) important in early steps of autophagosome biogenesis (Noda et al., 2000; Young et al., 2006). Aberrant autophagosome formation has been observed in cellular models of AP-4 deficiency (Davies et al., 2018; Mattera et al., 2017) as well as in the axons of neurons in *AP4B1* (Matsuda et al., 2008) and *AP4E1* (Ivankovic et al., 2020) knockout mouse models. The RUSC accessory proteins coordinate anterograde transport of ATG9A, SERINC1/3, and DAGLB toward the cell periphery in AP-4-derived vesicles (Davies et al., 2018, 2022). Conversely, Hook1 and Hook2 act as part of the FHF (FTS, Hook, and FHIP) complex thought to mediate retrograde trafficking of AP-4-coated and ATG9A-containing vesicles. This latter interaction is proposed to maintain a functional distribution of these proteins throughout the cytoplasm (Mattera et al., 2020). Despite being the first identified AP-4 accessory protein (Borner et al., 2012), the role of tepsin within the AP-4 coat has remained unclear.

Tepsin is a member of the epsin family of adaptor proteins (Borner et al., 2012), but structural and evolutionary evidence indicate it has functionally diverged (Archuleta et al., 2017). The tepsin unstructured C-terminus contains two conserved motifs for binding AP-4 appendage domains (Figure 1A; Frazier et al., 2016; Mattera et al., 2015). Tepsin contains two structured domains: an epsin N-terminal homology (ENTH) and VHS/ENTH-like (VHS-like) domains (Figure 1A; Frazier et al., 2016; Archuleta et al., 2017). X-ray crystallography structures of both domains revealed they lack critical features observed in other epsins that promote either phosphoinositide binding to ENTH domains or ubiquitin binding to VHS domains (Archuleta et al., 2017; Zouhar and Sauer, 2014). Tepsin has yet to be directly implicated in phenotypes associated with deficiencies in AP-4- trafficking or autophagy.

**Figure 1:**
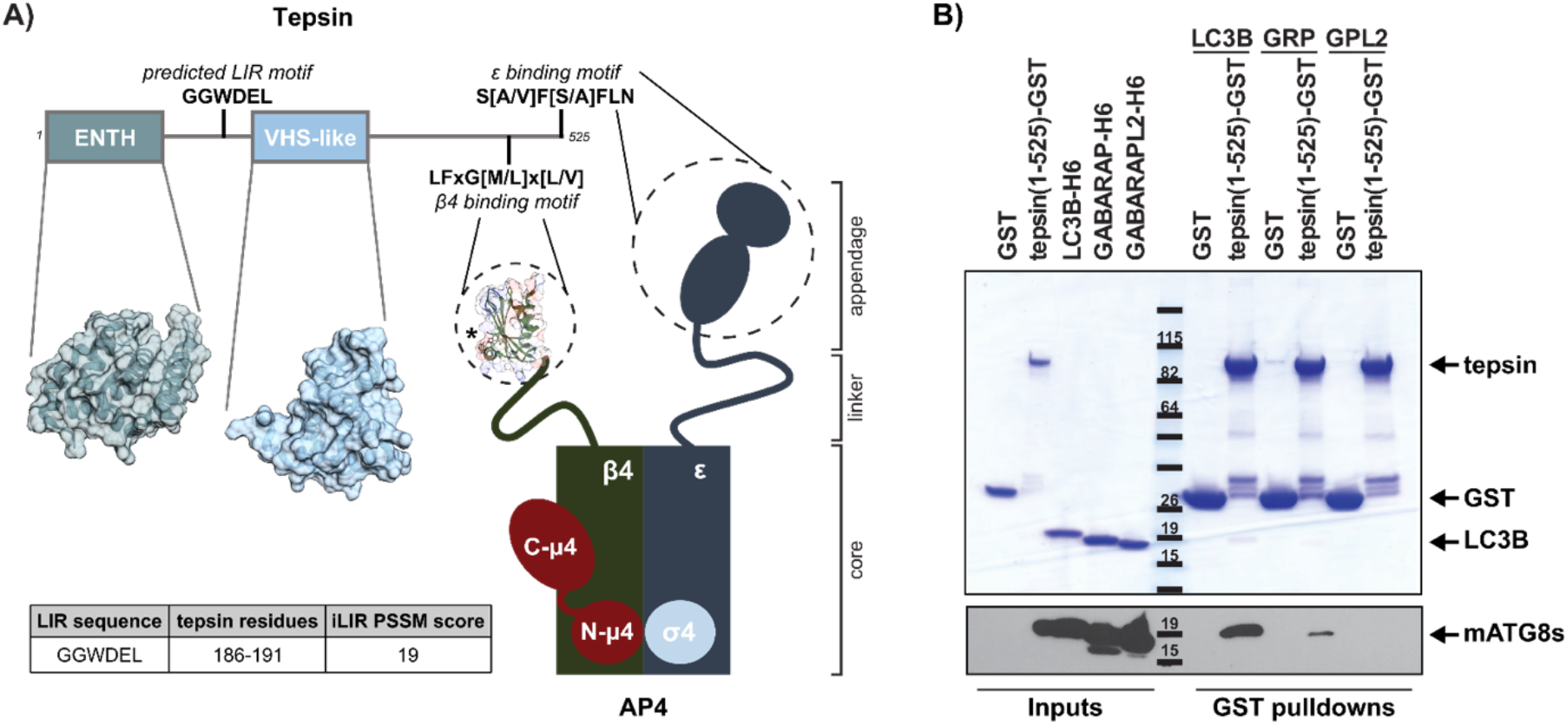
Tepsin directly and specifically binds LC3B *in vitro*. (A) Schematic diagrams of tepsin and AP-4 depicting the structural basis for the AP-4/tepsin interaction (PDB: 2MJ7); the predicted LIR motif (iLIR database; Jacomin et al., 2016) lies in the unstructured region between the tepsin ENTH (PDB: 5WF9) and VHS-like domains (PDB: 5WF2). (B) Coomassie-stained SDS-PAGE gel and Western blot (α-His; Abcam ab184607) of GST pulldowns using recombinant full-length tepsin-GST (residues 1-525) with His6x-LC3B, His6x-GABARAP (GRP), or His6x-GABARAPL2 (GPL2). Experiments show tepsin binds LC3B and weakly binds GABARAP; free GST was used as a negative control. Representative of three independent experiments.

Macroautophagy (hereafter autophagy) regulates cellular homeostasis by engulfing cytosolic material and dysfunctional organelles into autophagosomes that fuse with lysosomes for degradation and to promote macromolecule recycling within cells.

Many essential autophagy genes are conserved from yeast to humans, though higher eukaryotes possess an expanded number of autophagy-related proteins. This is exemplified by a key yeast autophagy protein, ATG8 (Feng et al., 2014). Mammals contain multiple ATG8 orthologs referred to as the mammalian ATG8 (mATG8) family. These proteins are further divided into two subfamilies: the LC3 (LC3A/B/C) and GABARAP (GABARAP and GABARAPL1, and GABARAPL2; Shpilka et al., 2011) families. The independent function of each mATG8 protein is not completely understood. They appear to have distinct roles at various stages of autophagy, from phagophore expansion and recruitment of autophagosome contents to maturation and closure of the autophagosome membrane (Lee and Lee, 2016). Thus far, biochemical analysis of both yeast and mammalian ATG8 proteins reveals many protein-protein interactions are mediated through short peptide motifs, commonly termed LIR motifs (<u>L</u>C3 Interacting <u>R</u>egion) in mammals (Birgisdottir et al., 2013).

In this study, we have identified a role for tepsin in regulating autophagy dynamics beyond export of ATG9A from the TGN. We demonstrate that tepsin directly interacts with LC3B *in vitro* and in cultured cell lines. The tepsin/LC3 interaction occurs between a LIR motif in tepsin’s first unstructured region (Figure 1A) and the LIR docking site on the surface of LC3B. Biochemical data and AlphaFold modeling suggest how the tepsin LIR motif conveys specificity for LC3 over other mATG8 proteins. Tepsin depletion in cells drives partial accumulation of ATG9A near the TGN and promotes accumulation of ATG9A at the cell periphery. Tepsin depletion also dysregulates the morphology of autophagosomes and autolysosomes. The effect of tepsin depletion partly differs from AP-4 depletion providing evidence for a distinct function for tepsin in AP-4 coated vesicles. Finally, we tested the functional relevance of the tepsin LIR motif in ATG9A trafficking. While wild-type tepsin can partially rescue ATG9A trafficking defects, tepsin constructs lacking the LIR motif cannot rescue ATG9A subcellular distribution to the wild-type pattern. Together, these data suggest tepsin is important for ATG9A-containing AP-4 vesicle formation and the tepsin/LC3B interaction contributes to the distribution of ATG9A in cells.

## Results

### *Tepsin directly and specifically binds LC3B* in vitro

The *in silico* iLIR database (Jacomin et al., 2016) identified a strongly predicted LIR motif in tepsin (Figure 1A). LIR motifs commonly contain the sequence [W/F/Y]xx[V/I/L], where critical hydrophobic residues are buried into corresponding hydrophobic pockets on the surface of mATG8 proteins (Popelka and Klionsky, 2015). The putative tepsin LIR motif fits the canonical sequence (GGWDEL). Two glycine residues immediately preceding the WDEL sequence in tepsin were also highlighted in the iLIR result, based on common sequences found in annotated functional LIR motifs (Jacomin et al., 2016). As is common for LIR motifs (Popelka and Klionsky, 2015), this motif lies in an unstructured region located between the tepsin ENTH and VHS-like domains (Figure 1A). The mATG8 subfamilies are partially distinguished by varying preferences for the specific sequence of LIR motifs (Rogov et al.; Wirth et al., 2019). We tested whether tepsin could bind three distinct mATG8 members. We purified full length tepsin (residues 1-525) as a Glutathione S-Transferase (GST)-fusion protein. GST pulldown assays using recombinant purified tepsin-GST with His-tagged LC3B, GABARAP, or GABARAPL2 show tepsin preferentially binds to LC3B *in vitro* (Figure 1B). The tepsin/LC3 interaction is faintly visible by Coomassie staining on an SDS-PAGE gel and further confirmed by probing 6x-histidine-tags on mATG8 proteins by Western blot. Tepsin-GST weakly bound GABARAP and showed no detectable binding at either Coomassie or Western blot level to GABARAPL2 *in vitro*.

Based on biochemical and cell biological data, tepsin requires AP-4 binding to become membrane-associated (Archuleta et al., 2017; Frazier et al., 2016; Borner et al., 2012; Mattera et al., 2015). Like yeast ATG8, all mATG8 proteins are conjugated to phosphatidylethanolamine through a ubiquitination-like cascade upon induction of autophagy (Shpilka et al., 2011). With this lipid modification, ATG8 proteins can be incorporated into the forming phagophore membrane to regulate autophagy. We tested whether AP-4 binding to tepsin affects the ability of tepsin to interact with LC3B using a tripartite GST pulldown assay. Tepsin-GST binding to the AP-4 β4 appendage domain does not significantly alter tepsin binding to LC3B *in vitro* (Figure S1A and S1B).

We next tested whether tepsin interacts with LC3B in cultured cells. Using an established HeLa cell line stably expressing tepsin-GFP (Borner et al., 2012), co-immunoprecipitation experiments show enrichment for endogenous LC3 (Figure S1C). Together, these data suggest an interaction between tepsin and LC3B occurs in cells, while the *in vitro* data using purified proteins suggests this is a direct interaction.

### The tepsin LIR motif binds the hydrophobic pocket on LC3

The critical aromatic residues in a LIR motif interact with LC3B using two hydrophobic pockets termed the LIR docking site (LDS). Using AlphaFold Multimer, we modeled binding between LC3B and a peptide containing the tepsin LIR motif (residues 185-193). The tepsin/LIR motif interaction was superposed over an experimentally determined structure of LC3B bound to a p62 LIR motif (PDB: 2ZJD). As observed in p62, the critical tryptophan and leucine residues are positioned in the hydrophobic pockets of the LC3 LDS (Figure 2A). Additionally, the LDS region contains corresponding basic patches to accommodate the acidic aspartate and glutamate residues of the tepsin LIR motif (W<u>DE</u>L).

**Figure 2:**
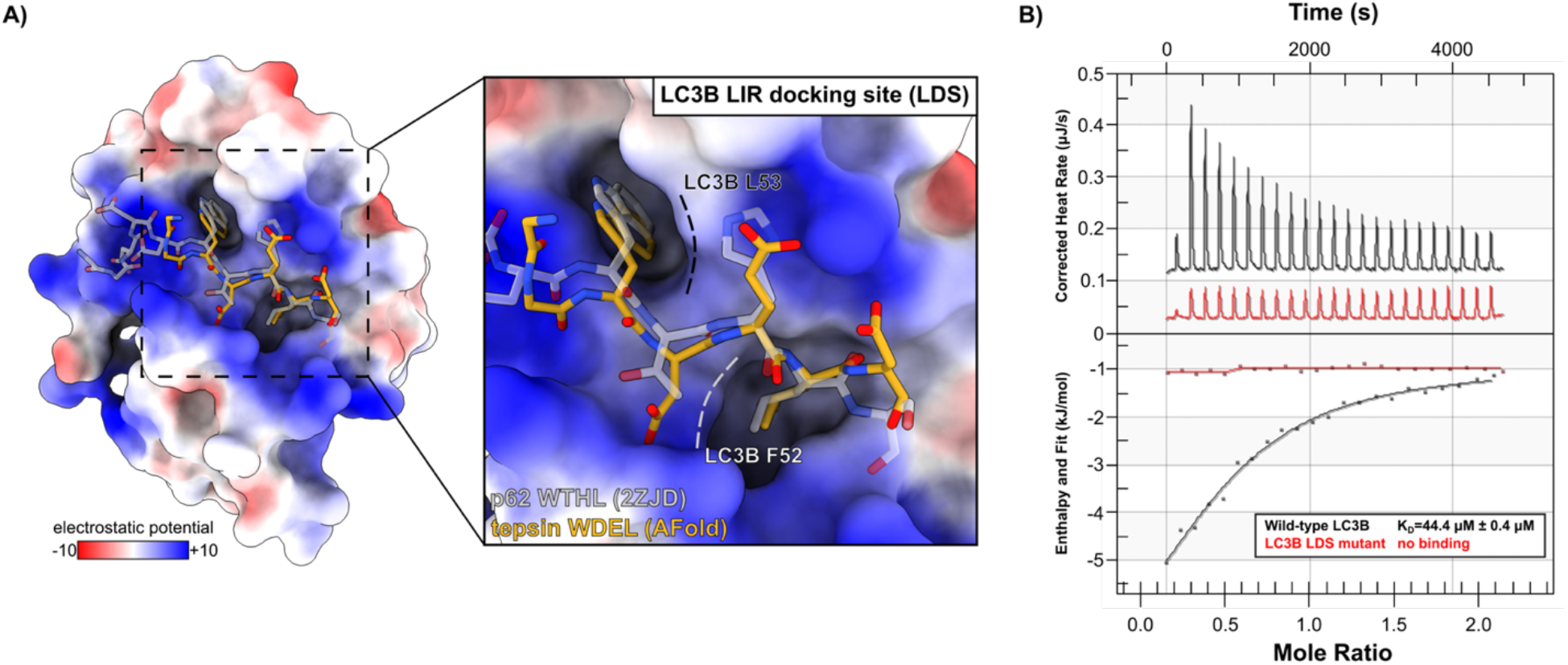
**The tepsin LIR motif binds the established binding pocket on LC3B *in vitro***. (A) Electrostatic surface representation of LC3B (PDB: 2ZJD) indicating the LIR Docking Site (LDS; shown in inset) bound to the LIR peptide in p62. A model generated in Alphafold Multimer shows the tepsin LIR motif peptide superposed (yellow) with the Trp and Leu residues occupying established hydrophobic pockets on the LC3B surface. (B) Representative isothermal titration calorimetry (ITC) experiments. Purified recombinant LC3B proteins and tepsin LIR motif peptide were used in ITC experiments to quantify binding affinities. The tepsin LIR motif peptide (residues 185-193) binds wild- type LC3B with a K_D_ of 44.4 µΜ ± 0.4 µΜ by ITC (n= 3 independent experiments). The tepsin LIR peptide shows no detectable binding to the LC3B LDS binding mutant (F52A/L53A). K_D_ values are represented as average ± standard deviation. ITC data are summarized in Table 1.

**Table 1:**
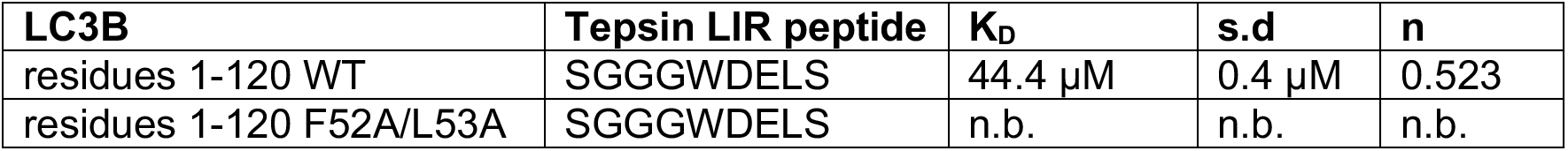
ITC data summary. This table summarizes representative ITC experiments from Figure 2, including protein constructs, relevant mutations, and peptide sequences; calculated K_D_ values; standard deviation (s.d.) and stoichiometry (n) values. “WT” denotes wild-type sequence; n.b. denotes no detectable binding.

| LC3B | Tepsin LIR peptide | K<sub>D</sub> | s.d | n |
| --- | --- | --- | --- | --- |
| residues 1-120 WT | SGGGWDELS | 44.4 μM | 0.4 μM | 0.523 |
| residues 1-120 F52A/L53A | SGGGWDELS | n.b. | n.b. | n.b. |

We used isothermal titration calorimetry (ITC) to quantify tepsin LIR motif binding to LC3B *in vitro*. Purified recombinant LC3B (residues 1-120) binds a synthesized tepsin LIR peptide (S<u>GGGWDELDS</u>, underlined residues 185-193) with micromolar affinity (*K_D_* = 44.4 μΜ ± 0.4 μΜ). Mutant LC3B containing point mutations in the LDS (F52A/L53A; Behrends et al., 2010; Marshall et al., 2019; Noda et al., 2008) exhibits no detectable binding to the tepsin LIR peptide (*K_D_* > 300 μΜ; Figure 2B). In the context of full length recombinant tepsin, pulldowns reveal mutation of either the tepsin LIR motif or the LC3B LDS abrogates binding at both Coomassie and Western blot detection levels (Figure S2A). Together, these biochemical data indicate the tepsin LIR motif mediates an interaction between tepsin and LC3B in the established LIR binding site (Noda et al., 2008).

To better understand the selectivity of the tepsin LIR motif for LC3B over other mATG8 proteins (Figure 1B), we utilized AlphaFold to generate models of tepsin LIR peptides bound to LC3A, GABARAP, and GABARAPL2. These models were then superposed with experimentally determined structures of each mATG8 bound to a LIR motif. Models of LC3A show tepsin LIR residues W188 and L191 fit well into the corresponding hydrophobic pockets (Figure S2B), as observed for LC3B models. The LC3A LDS also contains basic patches that could accommodate the acidic Asp and Glu residues in tepsin. The GABARAP model shows tepsin LIR residues W188 and L191 dock well into hydrophobic pockets (Figure S2C). The reduced binding potential observed biochemically for GABARAP may instead be explained by two acidic residues (D189 and E190) since the region between the hydrophobic pockets has a greatly reduced electrostatic potential compared to LC3A and LC3B. Finally, the GABARAPL2 model clearly exhibits a clash with the critical tepsin LIR W188 residue (Figure S2D).

Together these models support our biochemical data (Figure 1B) indicating the tepsin LIR motif selectively binds LC3 over GABARAP proteins.

### ATG9A accumulates near the periphery in cells acutely depleted of tepsin

We next sought to test the impact of tepsin knockdown in the context of established AP-4-deficiency phenotypes. Given the emerging links between AP-4 trafficking and autophagy (Mattera et al., 2020, 2017; Guardia et al., 2021; de Pace et al., 2018; Davies et al., 2018; Ivankovic et al., 2020; Matsuda et al., 2008), we utilized a well-established autophagy reporter line overexpressing a mRFP-GFP-LC3B fusion construct (Kimura et al., 2007; Sarkar et al., 2009). We acutely depleted these cells for tepsin or AP-4 using siRNA-mediated knockdown (Figure S3B). Whole lysate characterization by Western blot of these cells showed tepsin-depletion increased ATG9A expression by 1.5-fold (Figure S4A).

We tested whether and how the subcellular distribution of ATG9A might be altered upon tepsin knockdown by co-immunostaining for ATG9A and the *trans*-Golgi marker, TGN46 (Figure 3A). Tepsin-depleted cells exhibit decreased co-localization of ATG9A with the TGN marker as quantified by Manders’ overlap coefficient (Figure 3C). To further analyze this change to ATG9A subcellular localization, cells were segmented using the TGN46 marker into *trans*-Golgi ATG9A and non-Golgi ATG9A signal (representative enlargements in Figure 3B). In tepsin-depleted cells, isolated instances of ATG9A accumulation at the TGN were observed (Figure 3B.1; Figure 3D), but ATG9A largely redistributed to the cell periphery as judged by enlarged accumulations with high fluorescence intensity (Figure 3B.2; Figure 3E and 3F). Some peripheral accumulations of ATG9A were also observed upon AP-4 knockdown (Figure 3B.2).

**Figure 3:**
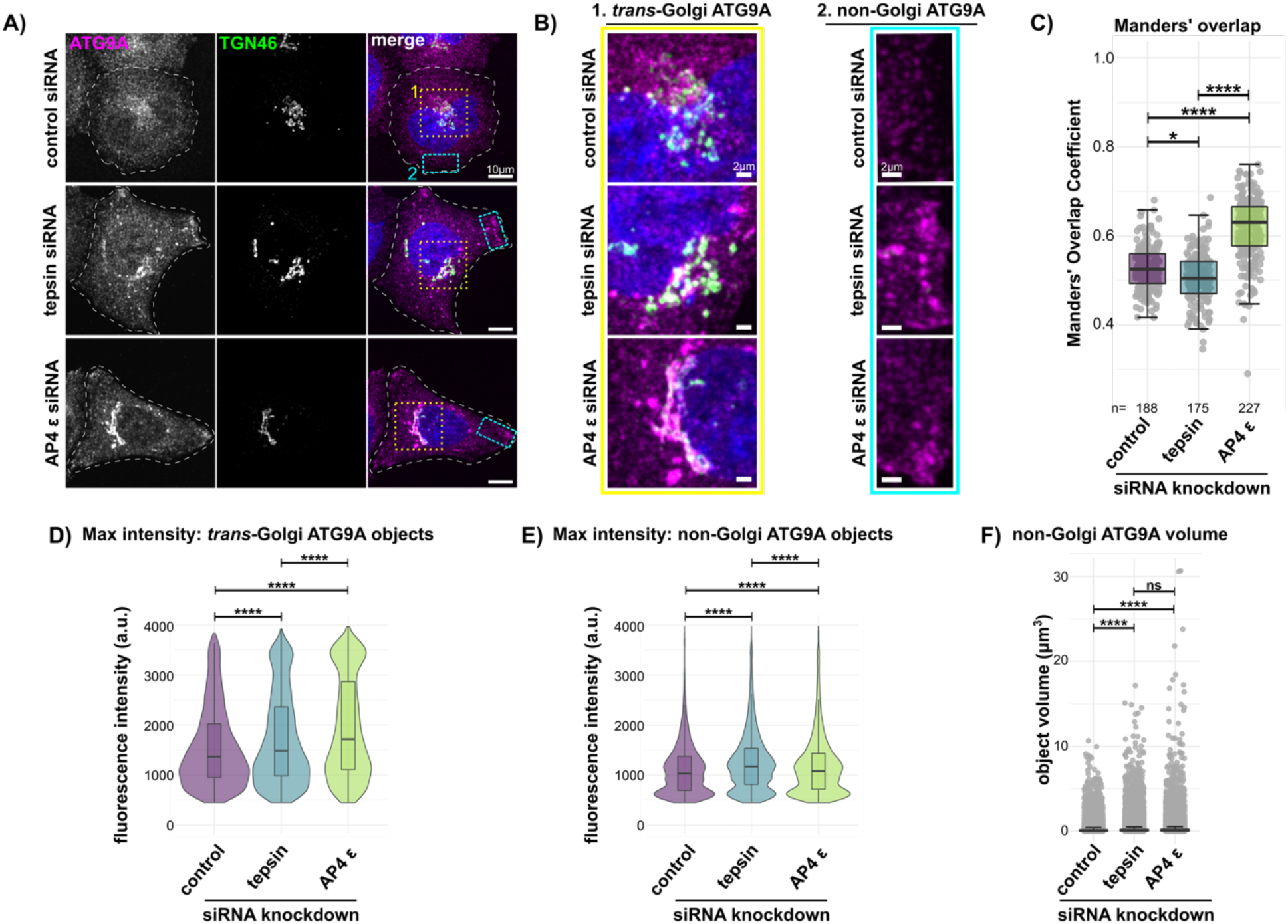
Tepsin depletion drives ATG9A accumulation at the cell periphery. (A) Representative maximum intensity projection confocal images of quenched mRFP- GFP-LC3B HeLa cells immunostained for ATG9A (α-ATG9A Abcam ab108338; secondary α-Rabbit-647 Thermo Fisher Scientific A32733) and TGN46 (α-TGN46 Bio- Rad AHP500GT; secondary α-Sheep-488 Thermo Fisher Scientific A11015). Cells were treated with control, non-targeting siRNA or tepsin siRNA. Scale bar: 10 µm. (B) Tepsin depletion results in ATG9A accumulation near the *trans*-Golgi (B.1; yellow boxes, A) and at the cell periphery (B.2; cyan boxes, A); Scale bar: 2 µm. (C) Manders’ overlap coefficient indicates on a whole cell level, less ATG9A co-localizes with TGN46 after tepsin depletion. ATG9A is enriched in the TGN in AP-4-depleted cells. (D-F) Quantification of high intensity ATG9A objects (details in Methods). (D) Tepsin-depleted cells exhibit increased intensity of ATG9A signal accumulated at the TGN, at lower levels than those observed in AP-4-depleted cells. (E) The maximum intensity of non- Golgi ATG9A objects is significantly increased in tepsin-depleted cells compared to both control and AP-4-depleted cells. (F) AP-4- or tepsin-depleted cells exhibit enlarged ATG9A object volumes (µm^3^). Quantification of four independent experiments (n=total cell count); each dot represents an individual cell (C) or ATG9A object (D-F). Statistical results from Kruskal-Wallis test, Dunn test with Bonferroni correction; ns>0.05, *p≤0.05, **p≤0.01, ***p≤0.001, ****p≤0.0001.

However, fluorescence intensity measurements indicate tepsin-depleted cells exhibit more apparent ATG9A in these peripheral structures than do AP-4-depleted cells (Figure 3E). The majority of ATG9A remains trapped in the TGN following AP-4 knockdown (Figure 3C, S4B, and S4C), as observed in multiple systems (Mattera et al., 2017; Davies et al., 2018; Ivankovic et al., 2020; Behne et al., 2020; Scarrott et al., 2023; de Pace et al., 2018). These data implicate tepsin in the ATG9A trafficking and export at the TGN, although it remains possible tepsin may also function in targeting these ATG9A-positive vesicles to their destination (next section; Discussion).

### Tepsin-depleted cells exhibit dysregulated autophagosomes and autolysosomes

The biochemical and cellular data presented thus far suggest there are additional functions for the AP-4 coat in regulating autophagy dynamics via the accessory protein, tepsin. We next used the mRFP-GFP-LC3B cell line to assay flux through the autophagic pathway (Figure 4A). Coincident red and green structures positive for mRFP-GFP-LC3 are identified as yellow autophagosomes (or early autophagic structures). Lysosome fusion with autophagosomes quenches GFP fluorescence, so red (mRFP-only) LC3 structures are classified as autolysosomes. We visualized autophagosome and autolysosome structures in tepsin-depleted or AP-4-depleted cells under basal or starvation (amino acid- and serum-free) conditions (Figure 4B).

**Figure 4:**
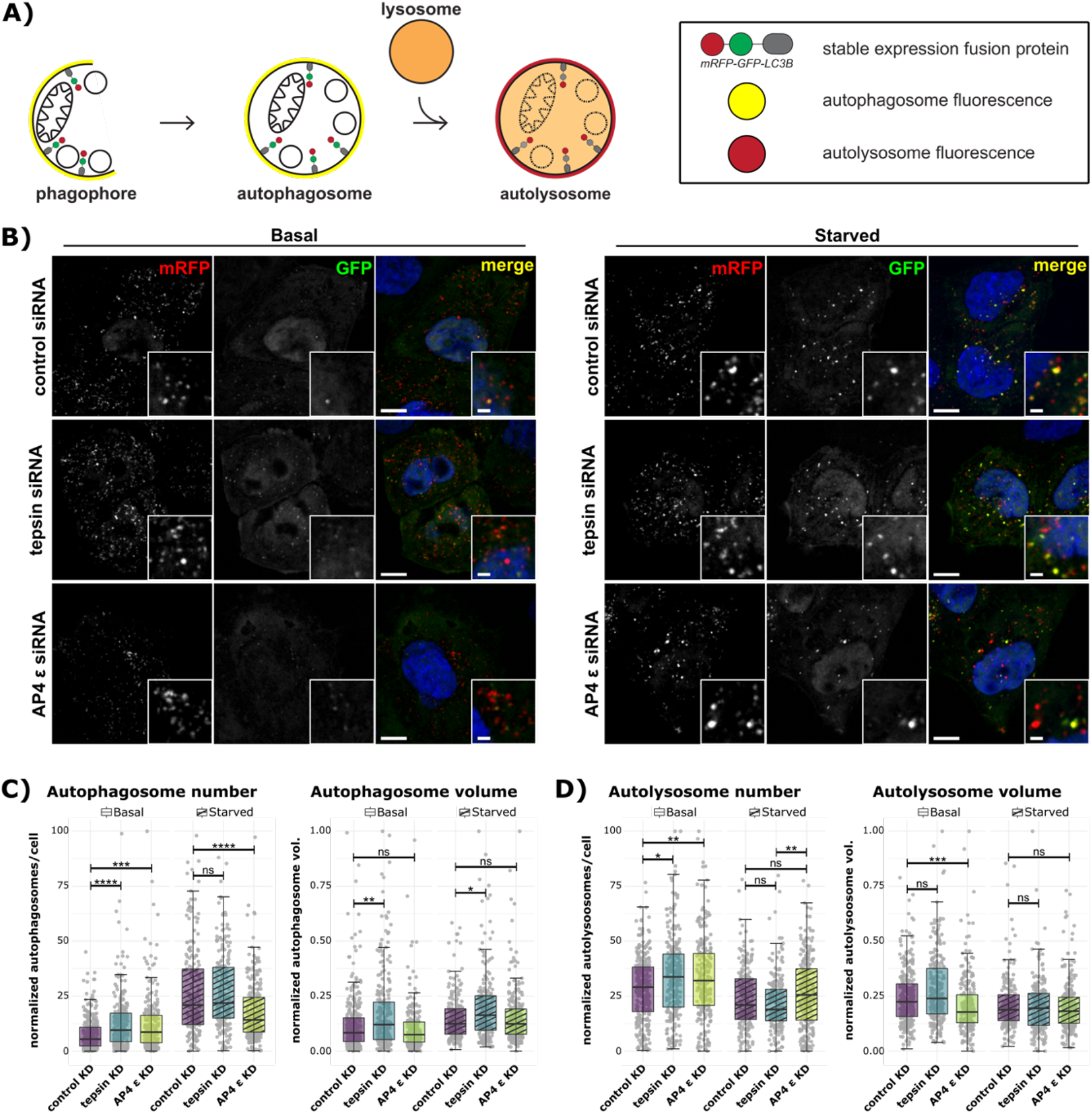
Acute tepsin depletion increases autophagosome number and volume under basal nutrient conditions. (A) Schematic of autophagy assay in HeLa cells stably expressing mRFP-GFP-LC3B. Coincident green and red fluorescence marks autophagosomes in yellow, while acidification of autolysosomes quenches GFP and leaves only red (RFP) signal. (B) Representative single plane confocal images of mRFP-GFP-LC3B HeLa cells cultured in either complete media (basal) or starved for 2 hours in EBSS. Cells were treated with control (non-targeting) siRNA, tepsin siRNA, or AP-4 ε siRNA. Scale bar: 10 µm; inset scale bar: 2 µm. (C-D) Quantification of 3 independent experiments with at least 200 total cells per condition. (C) Tepsin-depleted cells under basal conditions contain more autophagosomes with greater apparent autophagosome volume (µm^3^) under both basal and starved conditions. AP-4 depleted cells contain more autophagosomes under basal conditions but fewer autophagosomes in starved conditions. (D) Tepsin-depleted cells under basal conditions contain more autolysosomes but no apparent difference in autolysosome volume. AP-4-depleted cells under basal and starved conditions contain increased autolysosome number but decreased apparent autolysosome volume only under basal conditions. Plots for C and D depict individual cell averages for each metric overlayed with box-and-whisker plots of the distribution where each point represents one cell. Statistical results from Kruskal- Wallis test, Dunn test with Bonferroni correction. All data was subject to min-max normalization; ns>0.05, *p≤0.05, **p≤0.01, ***p≤0.001, ****p≤0.0001.

Knockdown efficiency and autophagy induction were confirmed by Western blot (Figure S5A).

Under basal conditions, both AP-4- and tepsin-depleted cells contain more autophagosomes (Figure 4C). Following starvation and induction of bulk autophagy, the number of autophagosomes in tepsin-depleted cells does not differ from the control treatment. However, AP-4-depleted cells contain fewer autophagosomes compared to control cells (Figure 4C). The apparent volume of autophagosomes in tepsin-depleted cells was increased under both basal and starvation conditions but was not significantly different from control in AP-4-depleted cells (Figure 4C). These data suggest tepsin- depletion results in the aberrant formation and growth of autophagosomes, while AP-4 acute depletion primarily affects the formation of early autophagosomes.

Tepsin or AP-4 depletion results in a greater number of autolysosomes in cells under basal conditions (Figure 4D). Autophagic flux is not drastically altered by tepsin depletion: starved cells show no significant change to the number of autolysosomes, and neither basal nor starved cells show changes to the apparent volume of autolysosomes. AP-4-depleted cells contain more autolysosomes compared to tepsin- depleted cells, but the number is not significantly different from control cells. Apparent autolysosome volume is decreased in AP-4-depleted cells under basal conditions compared to control cells (Figure 4D).

We further utilized the GFP tag in this reporter line to assay for an interaction with endogenous tepsin. GFP immunoprecipitations from mRFP-GFP-LC3B cells are enriched for endogenous tepsin and the AP-4 ε subunit by Western blot (Figure S5B). These data further indicate that tepsin interacts with LC3B in this cell line, and AP-4 is also present. Overall tepsin loss dysregulates autophagosome and autolysosome morphology without blocking autophagic flux. The effect of acute tepsin depletion is distinct from acute AP-4 depletion, suggesting a separate, or an additional, function for tepsin in autophagosomes biogenesis.

### The tepsin/LC3 interaction modulates AP-4-dependent ATG9A trafficking

Our data indicate tepsin loss impairs ATG9A trafficking and dysregulates autophagosome and autolysosome morphology. ATG9A peripheral accumulation when tepsin is lost suggests tepsin may target ATG9A-containing AP-4 vesicles for use in autophagy. We generated an siRNA-resistant, myc-tagged construct of wild-type tepsin to test whether ATG9A accumulation was specific to tepsin loss. Tepsin-myc expression level was similar between control and tepsin-siRNA treatments confirming siRNA resistance (Figure S6, WT tepsin). We monitored the colocalization of ATG9A with the TGN46 marker in the quenched mRFP-GFP-LC3B cell line. Untransfected cells depleted for tepsin exhibited decreased ATG9A signal at the TGN (Figure 5A and 5C). However, re-introduction of wild-type tepsin rescues high-intensity ATG9A accumulations and restores ATG9A intracellular distribution (as measured by TGN46 co-localization) to levels similar to those observed in control cells (Figure 5B and 5D). These data indicate ATG9A mis-trafficking (Figure 3) was specific to the loss of tepsin.

**Figure 5:**
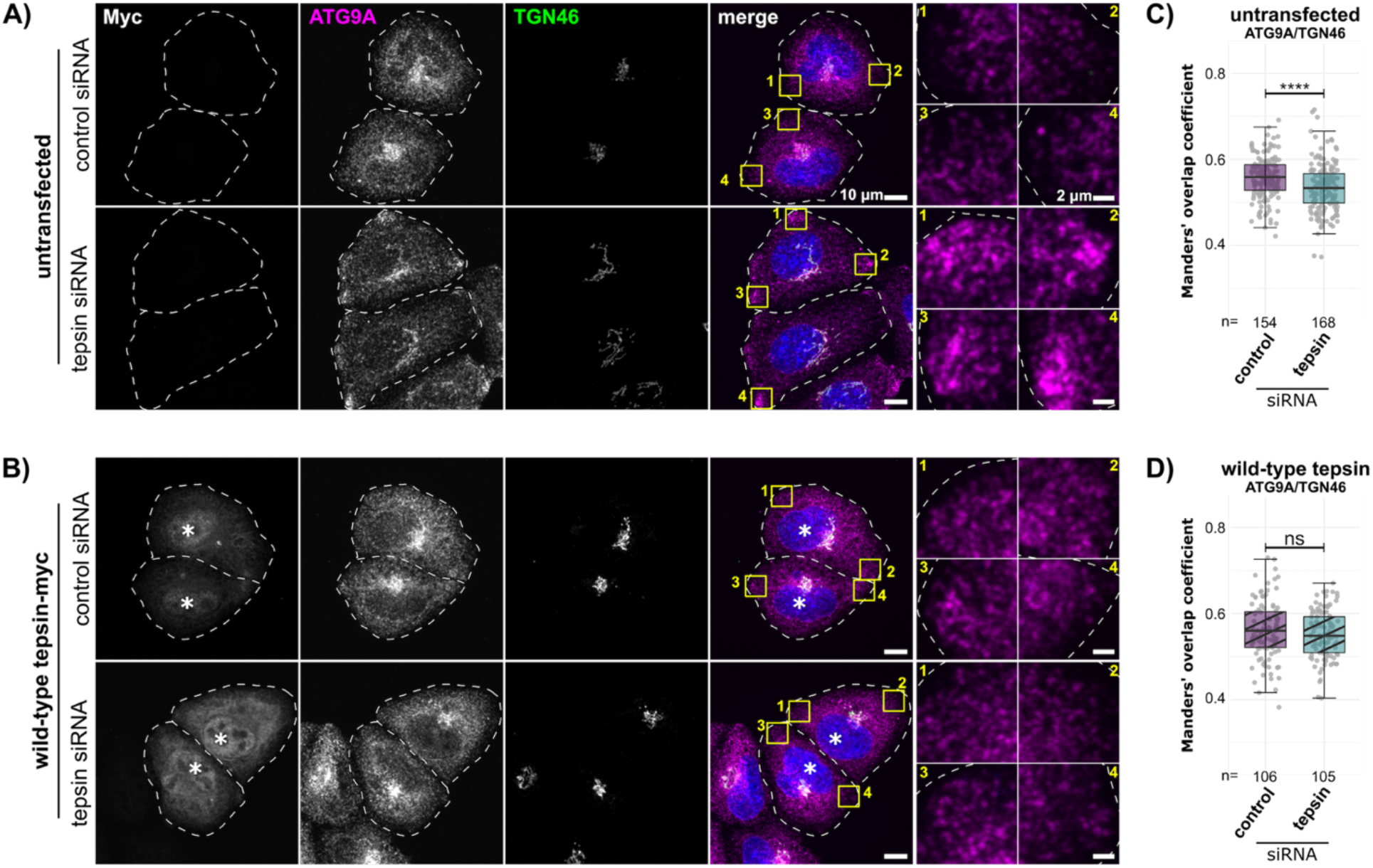
**Re-introduction of wild-type tepsin partially restores ATG9A cellular distribution in tepsin-depleted cells**. (A-B) Representative maximum intensity projection confocal images of quenched mRFP-GFP-LC3B HeLa cells immunostained for ATG9A (α-ATG9A Abcam ab108338; secondary α-Rabbit-647 Thermo Fisher Scientific A32733), TGN46 (α-TGN46 Bio-Rad AHP500GT; secondary α-Sheep-488 Thermo Fisher Scientific A11015), and myc-tag (α-myc-tag Cell Signaling Technology 2276; secondary goat α-Mouse-555 Thermo Fisher Scientific A32727). Cells were treated with non-targeting siRNA (control) or tepsin siRNA. Cells in B were subsequently transfected with wild-type siRNA resistant tepsin; transfected cells (anti- myc stained) are marked by asterisks (*). ATG9A accumulates at the cell periphery to a greater extent in tepsin-depleted cells (A; yellow inset boxes) compared to cells rescued with wild-type tepsin (B; yellow inset boxes). Scale bar: 10 µm; inset scale bar: 2 µm. (C) Manders’ overlap coefficient indicates less ATG9A resides within the *trans*-Golgi of tepsin-depleted cells. (D) ATG9A subcellular distribution is restored following re- introduction of wild-type tepsin. Quantification of 3 independent experiments (n = total cell number). Plots for C and D depict box-and-whisker plots of the distribution where each point represents one cell. Statistical results from Kruskal-Wallis test, Dunn test with Bonferroni correction; ns>0.05, *p≤0.05, **p≤0.01, ***p≤0.001, ****p≤0.0001.

Based on the *in vitro* characterization of the tepsin-LC3B interaction, we specifically tested the role of the tepsin LIR motif in ATG9A cargo trafficking. We generated siRNA-resistant myc-tagged constructs of LIR mutant tepsin (WDEL→SSSS) and a truncated tepsin containing only the C-terminal tail (ΔN-term tepsin; residues 360- 525). Each construct transfected at similar efficiency and exhibited resistance to tepsin siRNA treatments (Figure S6, ΔN-term and LIR mutant). Expression of LIR mutant tepsin failed to rescue ATG9A cellular distribution in tepsin-depleted cells (Figure 6A and 6C). Interestingly, instead of the TGN and peripheral accumulations previously observed, ATG9A is more diffusely distributed throughout the cell. We also tested the ΔN-term tepsin mutant, which contains only tepsin C-terminal tail harboring established AP-4 binding motifs (Frazier et al., 2016; Mattera et al., 2015). Expression of the ΔN- term tepsin mutant gives a dominant negative phenotype, in which ATG9A co- localization with the TGN46 marker is decreased in both control siRNA-treated and tepsin-depleted cells (Figure 6B and 6D).

**Figure 6:**
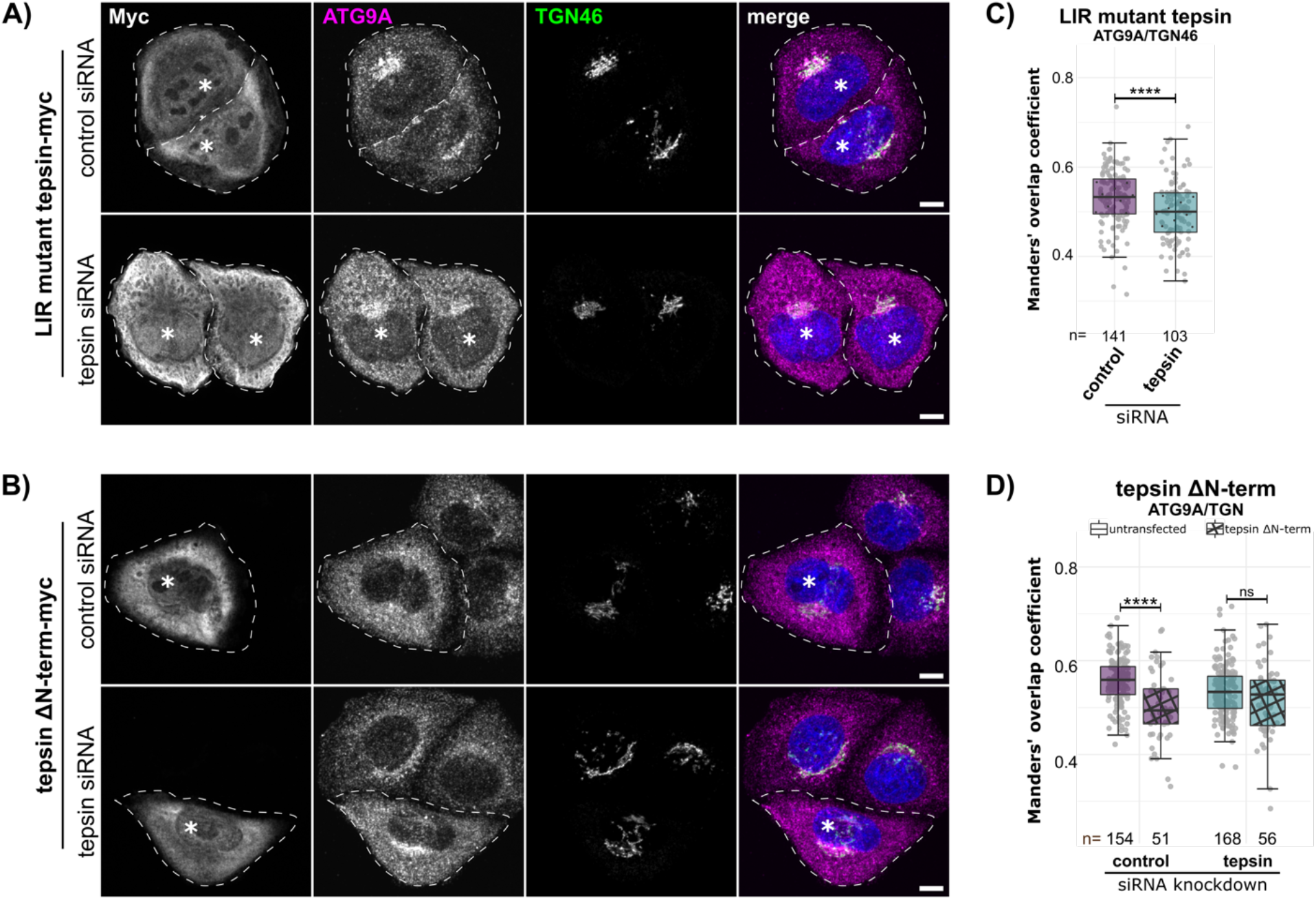
**The tepsin LIR motif is required to maintain ATG9A cellular distribution**. (A-B) Representative maximum intensity projection confocal images of quenched mRFP-GFP-LC3B HeLa cells immunostained for ATG9A (α-ATG9A Abcam ab108338; secondary α-Rabbit-647 Thermo Fisher Scientific A32733), TGN46 (α-TGN46 Bio-Rad AHP500GT; secondary α-Sheep-488 Thermo Fisher Scientific A11015), and myc-tag (α-myc-tag Cell Signaling Technology 2276; secondary goat α-Mouse-555 Thermo Fisher Scientific A32727). Cells were treated with non-targeting siRNA (control) or tepsin siRNA. Cells were subsequently transfected with LIR mutant tepsin (WDEL/SSSS) or tepsin ΔN-terminus (residues 360-525; ΔN-term); transfected cells are marked by asterisks (*). Targeted mutation of the LIR motif (A) or loss via N-terminus truncation (B) results in dispersed ATG9A accumulation outside of the *trans*-Golgi. Scale bar: 10 µm. (C) Introducing LIR-mutant tepsin in tepsin-depleted cells fails to restore ATG9A cellular distribution measured by Manders’ overlap coefficient for ATG9A signal at the TGN. (D) Tepsin ΔN-term acts as a dominant negative by dysregulating ATG9A colocalization at the TGN in control siRNA- and tepsin siRNA- treated cells. Quantification of 3 independent experiments (n = total cell number). Statistical results from Kruskal-Wallis test, Dunn test with Bonferroni correction in C and Mann-Whitney U-test in D; ns>0.05, *p≤0.05, **p≤0.01, ***p≤0.001, ****p≤0.0001.

Both the tepsin LIR mutant and ΔN-term tepsin constructs exhibited increased cytotoxicity (based on viability during sample preparation) and more variability in expression level for individual cells when visualized by immunofluorescence imaging. As a result, fluorescence intensity of both constructs is higher than for wild-type tepsin in many cells (represented in Figure 5A and 5B; 6A and 6B). However, for each construct, exogenous expression levels by Western blot were comparable between control and tepsin-depleted cells. Increased toxicity and dominant negative effects following ΔN-term tepsin transfection could indicate the tepsin N-terminus performs several important functions (see Discussion). Though both the LIR mutant and βN-term tepsin constructs retain AP-4 binding motifs, they fail to rescue ATG9A cellular distribution phenotypes. Together with the *in vitro* data, this suggests cells need a functional tepsin N-terminus and a direct interaction between tepsin and LC3B to mediate AP-4-dependent trafficking of ATG9A vesicles for their intended function during autophagy.

### Acute tepsin depletion reduces ATG9A and LC3 colocalization in cells

Following tepsin depletion, ATG9A distribution is disrupted and autophagosome morphology is altered in mRFP-GFP-LC3 HeLa cells. Since ATG9A distribution requires a functional tepsin LIR motif, we tested whether ATG9A accumulations coincide with LC3B structures in tepsin-depleted HeLa cells co-immunostained for endogenous LC3 and ATG9A (Figure S7A and S7B). Acute tepsin depletion (Figure S7C) results in decreased ATG9A/LC3 co-localization (Figure S7D and S7E), reflecting peripheral accumulations of ATG9A with no apparent LC3 signal (Figure S7A inset 2). In tepsin- depleted cells we also observed anomalous LC3 ring-like structures in a limited number of cells across independent replicates which could correspond to malformed or enlarged autophagosomes (Figure S7B). LC3-positive structures in control and tepsin-depleted cells typically coincide with ATG9A signal (Figure S7F) suggesting some ATG9A can still form maturing autophagosomes. Also supporting data from the mRFP-GFP-LC3B HeLa reporter cell line (Figure 4), endogenous LC3 structures in basal tepsin-depleted cells are larger (Figure S7G) and brighter (Figure S7H) than in control cells. These data further indicate tepsin contributes to proper ATG9A cellular distribution. Peripheral ATG9A accumulations do not colocalize with LC3; however, subpopulations of ATG9A can still contribute to autophagosome formation in tepsin-depleted cells.

## Discussion

### Summary

Published work from multiple groups has characterized the molecular interaction of tepsin with AP-4 (Frazier et al., 2016; Mattera et al., 2015; Borner et al., 2012) and determined X-ray structures of the tepsin ENTH and VHS-like domains (Archuleta et al., 2017). Despite these data, the function for tepsin in AP-4 coats has remained elusive. Data presented here provide multiple new mechanistic insights into tepsin function in AP-4-mediated trafficking. Using *in silico* and biochemical methods together with fluorescence imaging, we have identified and characterized a functional LIR motif in tepsin that interacts with LC3B *in vitro* and in cultured cells. Tepsin depletion in cells induces partial ATG9A accumulation at the TGN and enrichment at the cell periphery. Re-introduction of tepsin constructs with a mutated LIR motif or lacking the N-terminus are unable to rescue ATG9A trafficking defects. These data suggest tepsin may play roles in AP-4 vesicle formation and possibly in cargo delivery or recycling of ATG9A-positive vesicles (Figure 7).

**Figure 7:**
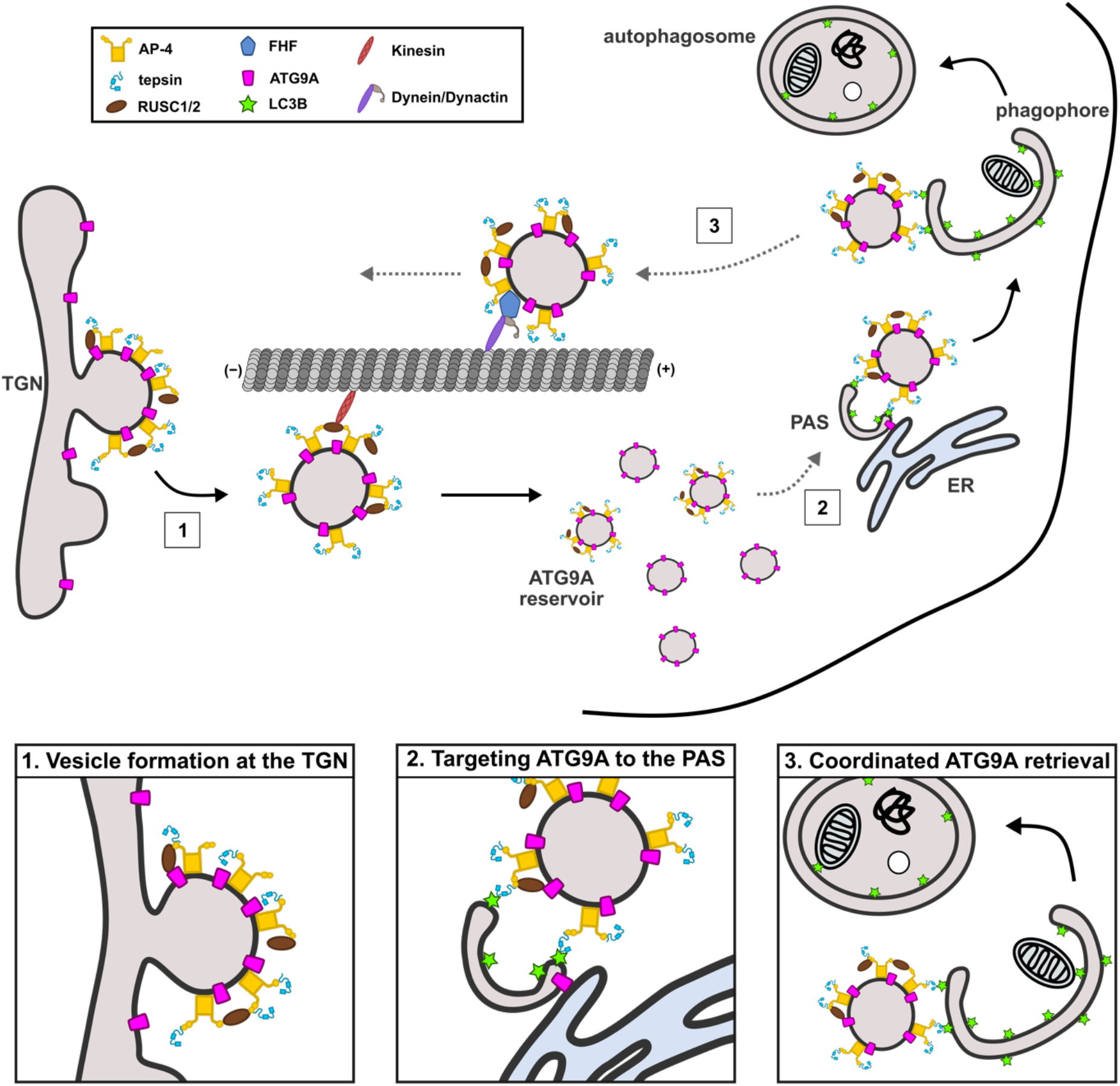
Models for tepsin function in AP-4 trafficking and autophagy. AP-4 coats containing tepsin assemble at the TGN, packaging ATG9A into vesicles. AP-4-coated and ATG9A-containing vesicles are proposed to undergo anterograde (Davies et al., 2018) and retrograde (Mattera et al., 2020) transport to maintain ATG9A distribution. Tepsin depletion leads to partial peripheral ATG9A accumulation which may suggest AP-4-coated vesicles function in ATG9A delivery. Together, published data and new data presented here lend themselves to three possible interpretations, that are not mutually exclusive. (1) Tepsin and AP-4 are required for efficient packaging of ATG9A into AP-4-coated vesicles exiting the TGN. (2) The tepsin/LC3B interaction may be required for targeting ATG9A to the PAS. (3) Alternatively, tepsin may bind LC3B to coordinate retrieval of ATG9A from maturing autophagosomes.

### Molecular characterization of the tepsin LIR motif

Emerging models of AP-4- deficiency syndrome highlight a dysregulation of autophagy resulting from retention of ATG9A, the essential autophagy lipid scramblase (Maeda et al., 2020; Guardia et al., 2020; Matoba et al., 2020; Noda et al., 2000; Young et al., 2006), at the TGN (Davies et al., 2018; Mattera et al., 2017; Ivankovic et al., 2020; Behne et al., 2020). Identification and characterization of a LIR motif within tepsin further links AP-4 coats to autophagy and maintenance of cellular homeostasis. The tepsin LIR motif (WDEL) fits an established consensus sequence (WxxL) for binding ATG8 proteins (Noda et al., 2008). Computational structural modeling and *in vitro* biochemical data provide strong evidence that tepsin interacts with LC3B with low micromolar affinity using the LIR docking site (LDS). LIR interaction affinities exhibit a large range (K_D_ values from sub-micromolar to ∼100 μM), usually because mATG8 family proteins exhibit selectivity (reviewed Wesch et al., 2020). This broad affinity range reflects the diverse functional roles of mATG8/LIR interactions and quantification of LIR motif binding affinities are not well explored (Birgisdottir et al., 2013).

The tepsin LIR motif shows partial specificity for LC3 over GABARAP proteins *in vitro* (Figure 1B). AlphaFold modelling indicates selectivity may be conferred by the tepsin acidic aspartate and glutamate residues (W<u>DE</u>L) which are better accommodated in the basic patch located between hydrophobic pockets in the LDS region of LC3 proteins (Figures 2 and S2). Furthermore, an alternative consensus sequence has been proposed as a GABARAP interaction motif (GIM), which displays a preference for aliphatic residues in the first variable position: [W/F]-[V/I]-X-V (Rogov et al., 2017). In cells, endogenous LC3B co-immunoprecipitates with tepsin-GFP (Figure S1C), and conversely, endogenous tepsin and AP-4 ε co-immunoprecipitate with mRFP-GFP- LC3B (Figure S5B). The micromolar affinity of this interaction *in vitro*, as well as weak detection by traditional immunoprecipitation methods, suggest the tepsin interaction with LC3B occurs transiently in cells.

The major question arising from the identification of the LIR motif is where in the cell do tepsin and LC3 interact? This question directly links to the major question in AP- 4 biology: where do AP-4 vesicles go? Both tepsin and LC3 cycle between cytosolic and membrane-associated states, with tepsin primarily localized at Golgi membranes during steady-state (Borner et al., 2012) while LC3 is found at autophagosomal membranes following lipid conjugation (Ichimura et al., 2000; Kabeya et al., 2000; Kirisako et al., 2000). LC3 is not enriched in AP-4 vesicle proteomic datasets which identified several AP-4 cargo and accessory proteins (Davies et al., 2018, 2022; Borner et al., 2012).

Proteomic vesicle profiling is imperfect for identifying prospective cargoes; however, LC3 lipid conjugation occurs via a ubiquitination-like cascade and seems to be targeted to sites of autophagosome biogenesis (Maruyama et al., 2021; Fracchiolla et al., 2016; Nath et al., 2014; Kabeya et al., 2004, 2000). It therefore seems unlikely LC3 is brought as a cargo by AP-4 vesicles, and instead supports the idea that tepsin transiently interacts with LC3B, likely at the autophagosomal membrane. Live cell imaging of HeLa cells stably expressing tepsin-GFP shows tepsin punctate structures form in the perinuclear region and persist to move peripherally within the cell before dissipating (Borner et al., 2012; Video 1).

LC3 conjugation begins early in autophagosome biogenesis; increases as the phagophore expands; and, following autophagosome closure, cytosolic-facing LC3 is deconjugated to its cytosolic form (Landajuela et al., 2016; Weidberg et al., 2010; Kirisako et al., 2000; Nair et al., 2012). Tepsin-LC3B interactions at phagophore membranes would likely be regulated by avidity effects dependent on protein density on the membrane. Autophagosome maturation and eventual deconjugation of LC3 could also temporally regulate the tepsin/LC3B interaction. It is also possible that other AP-4 coat components interact with autophagy machinery. The iLIR database (Jacomin et al., 2016) identifies multiple LIR motifs in another AP-4 accessory protein, RUSC2 (Davies et al., 2018). In future, it will be interesting to explore functionality of putative RUSC2 LIR motifs and whether they affect ATG9A trafficking.

### The tepsin/LC3B interaction affects ATG9A cellular distribution

Impaired export of ATG9A from the TGN is now established as a hallmark of AP-4-deficiency syndrome (Behne et al., 2020). AP-4 knockout in cultured cells (Davies et al., 2018; Mattera et al., 2017), fibroblasts from AP-4-deficient patients (Davies et al., 2018; de Pace et al., 2018), AP-4 ε or β4 knockout mice neurons (de Pace et al., 2018; Ivankovic et al., 2020; Scarrott et al., 2023), and the acute AP-4 depletion shown here (Figure 3C-D) robustly traps ATG9A at the TGN. Tepsin knockout HAP1 cells did not accumulate ATG9A at the TGN (Mattera et al., 2017), but the role of tepsin in ATG9A trafficking was largely unexplored. Data shown here, using acute depletion methods, offers compelling evidence that tepsin is required for efficient formation of AP-4 coated vesicles and/or delivery of ATG9A-positive vesicles.

Acute tepsin depletion has two distinct effects on ATG9A trafficking: it drives partial ATG9A accumulation near the TGN (Figure 3D) and promotes ATG9A accumulation at the cell periphery (Figure 3E-F). Accumulations at and near the TGN may reflect abortive or malformed AP-4 vesicles unable to leave the TGN vicinity. Re- introducing wild-type tepsin rescues both phenotypes (Figure 5). Whole cell lysates also exhibit increased ATG9A expression upon tepsin loss (Figure S4A). Similar ATG9A expression levels are increased in patient-derived cells (Davies et al., 2018) presumably as cells attempt to compensate for failed TGN export of ATG9A. In this work, acute tepsin-depletion led to slightly higher (1.5-fold increase) ATG9A expression levels than did AP-4-depletion (1.25-fold increase). Compensatory ATG9A overexpression in tepsin-depleted cells may be more effective, since ATG9A export is not blocked as significantly.

ATG9A has a well-documented role during autophagy, so we utilized an established mRFP-GFP-LC3B reporter (Sarkar et al., 2009) to assay flux through the autophagy pathway following acute depletion of tepsin or AP-4. Autophagic flux seemed largely unaffected by tepsin- or AP-4 depletion (Figure 4). However, formation and morphology of autophagosome and autolysosome structures were differentially affected by these depletions, suggesting some separation of function between tepsin and AP-4. We note AP-4 depleted cells form more autophagosomes under basal conditions but appear to have fewer autophagosomes than do control cells following autophagy induction (Figure 4C). This assay cannot distinguish between early phagophore structures and maturing autophagosomes, so this may indicate AP-4 loss hinders proper formation of early autophagic structures. ATG9A is critical for early nucleation of the phagophore (Young et al., 2006; Noda et al., 2000; Sawa-Makarska et al., 2020; Yamamoto et al., 2012), and AP-4 trafficking of ATG9A is thought to contribute to maintenance of an ATG9A vesicle pool used in these nucleation steps (Davies et al., 2018). Altered formation of autophagosomes without a total block in autophagy supports the idea that AP-4-derived ATG9A vesicles partially contribute to this pool.

Conversely, tepsin depletion primarily appears to affect morphology of autophagosomes which have larger apparent volume under basal or starved cell conditions (Figure 4C). A greater number of autophagosomes and autolysosomes form in tepsin-depleted cells, suggesting dysregulation of autophagy even under fed cell conditions (Figure 4C and 4D). If tepsin was solely involved in formation of AP-4 coated vesicles at the TGN, we would expect tepsin depletion to have the same effect on autophagy dynamics as AP-4-depletion because of failed ATG9A export. We observe slightly increased ATG9A expression in tepsin-depleted cells (Figure S4A), which could partially compensate for inefficient ATG9A export (Figure 7.1). The different effect of AP-4 or tepsin depletion on autophagosome morphology presented here (Figure 4) may reflect a secondary role for tepsin during autophagy.

The tepsin LIR motif binding to LC3B hints tepsin can mediate an interaction between AP-4 vesicles and autophagic membranes. Expressing tepsin with a mutated LIR motif failed to restore ATG9A subcellular distribution in tepsin-depleted cells. In fact, expression of the LIR mutant tepsin seems to drive diffuse distribution of ATG9A throughout the cell (Figure 6A and 6C). Similarly, expressing the ΔN-term tepsin fragment acts as a dominant negative, where ATG9A is expressed more diffusely throughout the cell in control cells (Figure 6B and 6D). The tepsin LIR mutant and ΔN- term retain established AP-4 binding motifs (Mattera et al., 2015; Frazier et al., 2016), so these results likely reflect specific contributions from the tepsin LIR motif and N-terminus. Additionally, ATG9A peripheral accumulations do not colocalize with LC3 (Figure S7).

ATG9A vesicles serve as the seed membrane for autophagosomes preceding the recruitment of additional autophagy machinery (Olivas et al., 2023; Broadbent et al., 2023). How and where ATG9A vesicles are coordinated for autophagy is not yet clear. These presented data suggest the tepsin LIR motif helps coordinate localization of ATG9A-positive vesicles to pre-autophagic sites (Figure 7.2). AP-4 trafficking is also implicated in retrograde transport of ATG9A (Mattera et al., 2020), so it remains possible that tepsin coordinates ATG9A retrieval (Figure 7.3). The recycler complex (sorting nexins 4, 5, and 17) mediates retrieval of some autophagy machinery from late autophagosome/autolysome membranes (Zhou et al., 2022). Our data do not exclusively favor a retrieval model as we would expect more ATG9A to coincide with LC3. For either of these models, it will be important for future work to establish whether AP-4 vesicles remain coated to mediate effects at the periphery. Live cell imaging in tepsin-GFP HeLa cells suggests AP-4 coated vesicles persist while moving away from the TGN toward the cell periphery (Borner et al., 2012; Video 1). Considering tepsin depletion results in morphological defects of early autophagosomes (Figure 4C and Figure S7G/H), the tepsin LIR interaction with LC3B may contribute to ATG9A function or delivery for development and growth of autophagy membrane structures. It may be particularly interesting to explore avidity effects on the tepsin-LC3B interaction given that maturation of autophagosome structures coincides with LC3 protein density at the membrane (Kabeya et al., 2000; Nath et al., 2014). Altogether these data identify the first functional role of tepsin and further develop our understanding of AP-4 trafficking and autophagy.

## Materials and Methods

### Reagents

Unless otherwise noted, all chemicals were purchased from Sigma.

The following antibodies were used in this study: rabbit anti-AP-4 epsilon 1:400 for Western blots (612019; BD Transduction Labs); rabbit anti-ATG9A 1:200 for immunofluorescence and 1:1000 for Western blots (ab108338; Abcam); rabbit anti- LC3B 1:3000 for Western blots (ab48394; Abcam); mouse anti-LC3 1:200 for immunofluorescence (M152-3; MBL); HRP-conjugated anti-GFP 1:2000 for Western blots (ab6663; Abcam); rabbit anti-tepsin 1:500 (in-house; Genscript) and 1:1000 (Robinson Lab, Cambridge) for Western blots; sheep anti-TGN46 1:1000 for immunofluorescence (AHP500GT; Bio-Rad); mouse anti-alpha tubulin 1:3000 for Western blots (66031; Proteintech); mouse anti-Myc-tag 1:8000 for immunofluorescence and 1:6000 for Western blots (2276; Cell Signaling Technology); HRP-conjugated anti-6X His tag 1:8000 for Western blots (ab184607; Abcam); HRP-conjugated secondaries for Western blots, 1:5000: Pierce goat anti-rabbit IgG (31460; Thermo Fisher Scientific); Pierce goat anti-mouse IgG (31430; Thermo Fisher Scientific); Fluorescent secondary antibodies for immunofluorescence, 1:500: goat anti-Rabbit 647 (A32733; Thermo Fisher Scientific), goat anti-Mouse 555 (A32727; Thermo Fisher Scientific), goat anti-Mouse 488 (A32723; Thermo Fisher Scientific), donkey anti-Sheep 488 (A11015; Thermo Fisher Scientific).

### Molecular biology and cloning

For biochemical analysis, full-length tepsin (Borner et al., 2012) was subcloned using *Nde*I/*Bam*HI sites into in-house vector pMW172 (Owen and Evans, 1998) modified to incorporate a C-terminal, thrombin cleavable GST tag. Full-length constructs of LC3B (residues 1-125), GABARAP (residues 1-117), and GABARAPL2 (residues 1- 117) were subcloned by Genscript into pGEX-6P-1 using *Bam*HI/*Sal*I sites to generate N-terminal GST-fusion protein; an additional His6x-tag was added to the N-terminus of each protein sequence. The GST-fusion AP-4 β4 appendage (residues 615-739) was cloned previously (Frazier et al., 2016). A two-stage mutagenesis protocol (Frazier et al., 2016) was used to truncate full-length LC3B into the mature isoform (residues 1- 120) and make the following mutations to tepsin and LC3B constructs: LC3B LIR docking site (LDS) mutant (F52A/L53A); LIR mutant tepsin (W188S, D189S, E190S, L191S); and tepsin siRNA target site silent mutations (nucleotides G1329A, A1335G, T1338C, A1341G, G1344A).

To generate constructs for tepsin rescue experiments, full-length tepsin was subcloned into pcDNA3.1 vector (Thermo Fisher Scientific) using *Bam*HI/*Xho*I sites. A C-terminal myc-tag (EQKLISEEDL) was included in the 3’ primer to generate tepsin- myc constructs. This construct was subjected to mutagenesis as described above to confer siRNA resistance and mutate the tepsin LIR motif. A truncated construct of the tepsin C-terminal tail (residues 360-525 with myc-tag) was subcloned from the siRNA-resistant tepsin-myc using *Bam*HI/*Sal*I sites into the pcDNA3.1 backbone. The sequences of all constructs described above were verified by Sanger DNA sequencing (Azenta). Oligonucleotides used in this study may be found in Table 2.

**Table 2:**
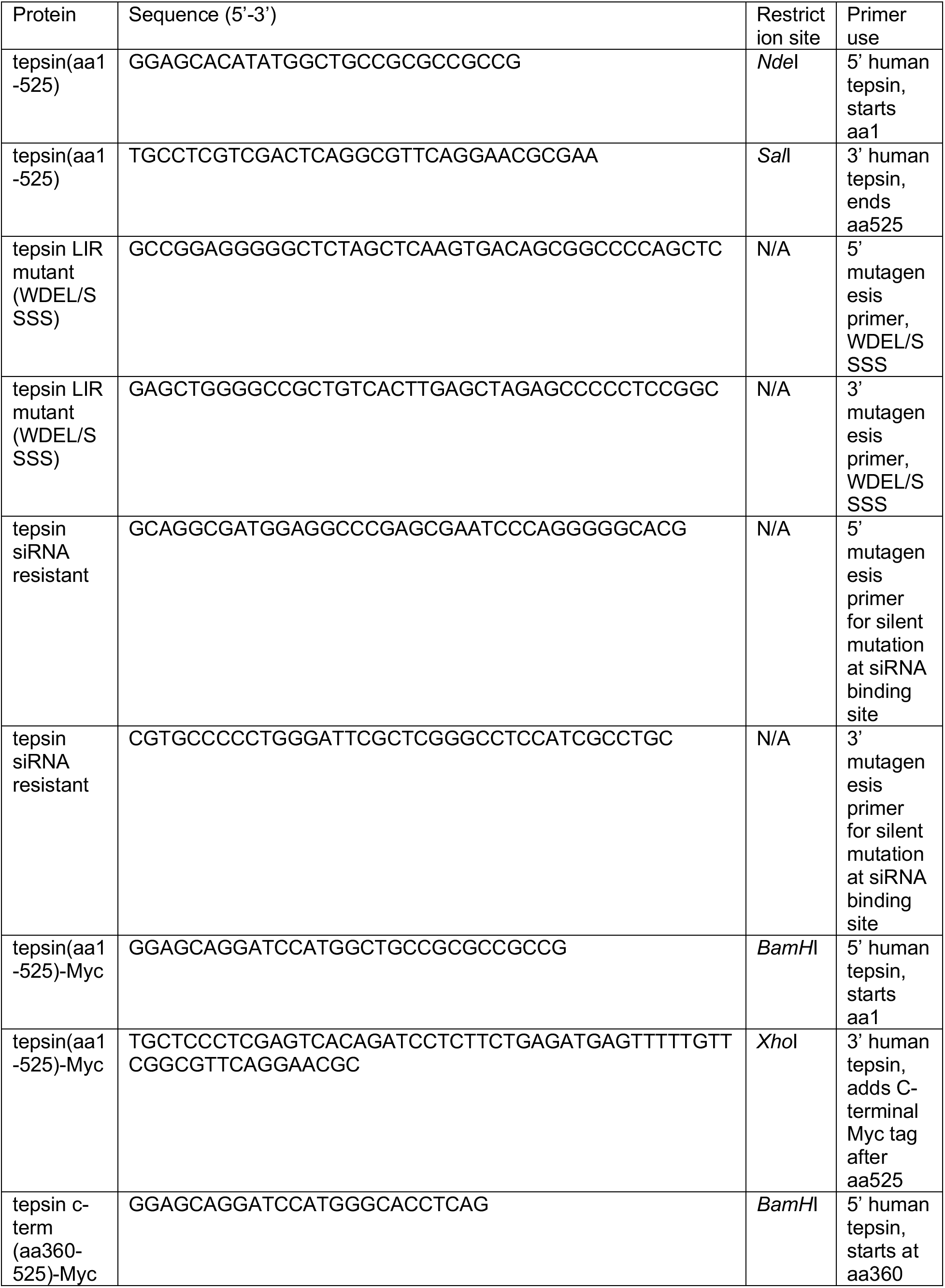
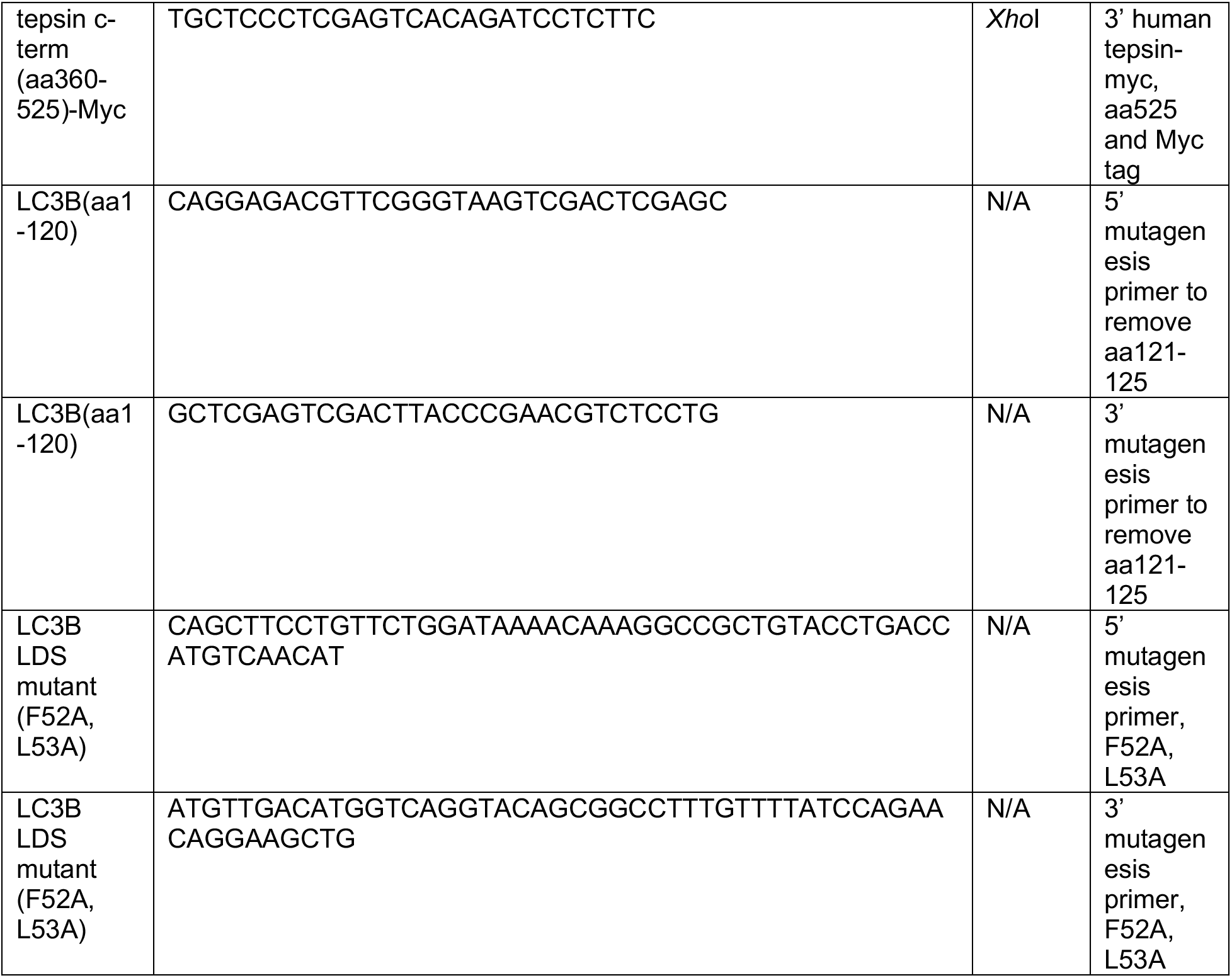
Oligonucleotides used in this study.

### Protein expression and purification

Constructs were expressed in BL21(DE3)pLysS cells (Invitrogen) for 16-20 hours at 22°C after induction with 0.4 mM Isopropyl β-D-1-thiogalactopyranoside (IPTG). All tepsin constructs were purified in 20 mM Tris (pH 8.5), 200 mM NaCl and 2 mM βME. Wild-type β4 appendage domain and all mammalian ATG8 proteins (mATG8; LC3B, GABARAP, and GABARAPL2) were purified in 20 mM HEPES (pH 7.5), 200 mM NaCl and 2mM βME. Cells were lysed by a disruptor (Constant Systems Limited) and proteins were affinity purified using glutathione sepharose (GE Healthcare) in purification buffer. GST-tagged mATG8 constructs were cleaved overnight with recombinant 3C protease at 4°C and eluted in batch. All proteins were further purified by gel filtration on preparative Superdex HiLoad 26/600 or analytical (Superdex 200 10/300) columns (GE Healthcare).

### Tepsin antibody production

The tepsin ENTH domain was expressed and purified as described previously (Archuleta et al., 2017). Protein was provided to Genscript to raise an antibody in New Zealand rabbits through three rounds of immunization followed by affinity purification. This tepsin antibody was tested on lysates generated from commercially available HAP1 AP-4 ε (gene: *AP4E1*) and tepsin (gene: *ENTHD2*) KO cell lines (Horizon Discovery) and compared to a published tepsin antibody (Figure S3A) provided by the Robinson lab (Borner et al., 2012).

### iLIR

Putative LIR motifs discussed in this paper were identified using the iLIR *in silico* resource (https://ilir.warwick.ac.uk/kwresult.php; Jacomin et al., 2016). The putative tepsin LIR motif is found in the list of identified putative LIR motif-containing proteins found searching by gene name (*ENTHD2*). It is also predicted when given the fasta sequence of full-length tepsin.

### Structure representation and modeling

Models of tepsin LIR motif interactions with mammalian ATG8 proteins were generated using AlphaFold 2.2 Multimer (Jumper et al., 2021b; a). Models were validated by visualization of peptide residues docked into hydrophobic pockets on the modeled mATG8 surfaces. Each model docked critical LIR motif residues (Trp188 and Leu191) into hydrophobic pockets without major clashes. The models were compared to experimental mATG8-LIR structures by superposition. ChimeraX MatchMaker was used to superpose Alphafold2 models with experimental structures deposited in the Protein Data Bank. All structural figures were generated in ChimeraX.

### GST-pulldown assays

GST or tepsin-GST fusion proteins (50 µg) and indicated prey proteins at a 5X molar excess to tepsin (mATG8 and/or AP-4 β4 appendage) were incubated with glutathione sepharose resin (GE Healthcare) for 2 hours at 4°C in 20 mM HEPES (pH 7.5), 100 mM NaCl, 0.5% NP-40 and 2 mM DTT. Resin was washed three times with the same buffer. Proteins were eluted from the resin using the wash buffer plus 30 mM reduced glutathione. Elution buffer was incubated with resin for 20 min on ice with gentle agitation every 2 minutes. Gel samples were prepared from the supernatant following elution with glutathione, and the assay was analyzed by Coomassie staining and Western blotting of SDS-PAGE gels. When further analyzed by Western blot, His-tagged prey protein were detected using anti-His6-HRP conjugated primary antibody (Abcam, ab184607). Uncropped gels and films are displayed in Figure S7.

### Isothermal titration calorimetry

Isothermal titration calorimetry (ITC) experiments were conducted on a NanoITC instrument (TA Instruments) at 25°C. Molar peptide concentration in the syringe was at least 6.5 times that of protein in the cell; 0.13 mM LC3B was placed in the cell and 0.845 mM tepsin LIR peptide was placed in the syringe. The tepsin LIR peptide was synthesized (Genscript) with the native tepsin sequence (residues 185-193; <u>S</u>GGGWDELS) and an additional serine added for solubility (underlined). All experiments were carried out in 20 mM HEPES (pH 7.5), 100mM NaCl and 0.5 mM TCEP, filtered and degassed. Incremental titrations were performed with an initial baseline of 120 seconds and injection intervals of 200 seconds. Titration data were analyzed in NANOANALYZE (TA Instruments) to obtain a fit and values for stoichiometry (n) and equilibrium association (K_a_). K_D_ values were calculated from the association constant.

### Tissue culture

HAP1 cell lines (Horizon Discovery; *AP4E1* KO: HZGHC000768c003; *ENTHD2* (tepsin) KO: HZGHC000845c002; Parental HAP1: C631) were maintained in IMDM (Gibco) supplemented with 10% v/v fetal bovine serum (FBS; R&D Systems). Stable tepsin-GFP (Borner et al., 2012) and mRFP-GFP-LC3B (Sarkar et al., 2009) HeLa cell lines were obtained as gifts from the Robinson and Rubinsztein labs (Cambridge Institute for Medical Research), respectively. Wild-type HeLa cells (ATCC) and stable HeLa cell lines were maintained in MEM-alpha (Gibco), supplemented with 10% v/v fetal bovine serum at 37°C in a 5% CO_2_ atmosphere. Media for stable cell lines was additionally supplemented with 600 µg/mL G418 (Corning). Cell lines were routinely monitored for mycoplasma contamination using DAPI to stain DNA. For starvation during autophagy assays, cells were washed twice with Earle’s balanced salt solution (EBSS; Gibco) and incubated with EBSS for 2 hours. Basal cells were concurrently given fresh complete media for the same duration. Where indicated, cells were treated with 100 nM Bafilomycin A1 (Millipore-Sigma) in either EBSS or complete cell media for 2 hours.

### Western blotting

Cells were resuspended and lysed in NP-40 lysis buffer (10 mM HEPES [pH 7.5], 150 mM NaCl, 0.5 mM ethylenediaminetetraacetic acid (EDTA), 1% NP-40, and 1 cOmplete Mini EDTA-free Protease Inhibitor Cocktail (Roche). Cell slurry was vortexed briefly then incubated on ice for 30 minutes. The cell slurry was then centrifuged at 20,500 x RCF for 30 minutes. The soluble fraction (lysate) was reserved, and total protein concentration was measured using Precision Red (Cytoskeleton). Normalized samples were denatured with SDS loading buffer (250mM Tris [pH 6.8], 50 mM DTT, 10% v/v SDS, 20% v/v glycerol, 0.5% w/v bromophenol blue) and boiled for 1 min at 95°C. Samples were subjected to SDS-PAGE using 4-20% gels (Bio-Rad) and transferred to PVDF membranes (Immobilon-P; Millipore). Blots were incubated with indicated primary and HRP-conjugated secondary antibodies then detected using Amersham ECL Western blotting reagents (GE Healthcare) or for more sensitive detection SuperSignal™ West Pico PLUS Chemiluminescent Substrate (Thermo Scientific. All uncropped blots are shown in Figures S8 and S9. Where indicated, Western blot results were quantified using the ImageStudio lite software (LI-COR).

Statistical significance was analyzed using R (R Core Team, 2021) with scripts written in RStudio to run one-way ANOVA with Tukey post-hoc test or Student’s t-tests as appropriate. The following packages were used in R for data management, statistical testing, and data visualization: tidyverse (Wickham et al., 2019); rstatix (Kassambara, 2021); and viridis (Garnier et al., 2021).

### GFP immunoprecipitation assays

Cell lysates from tepsin-GFP or mRFP-GFP-LC3B HeLa cells were prepared as described above. Lysate was incubated with unconjugated agarose (control) resin (Chromotek) for 30 min at 4°C to reduce background binding. This pre-cleared lysate was divided equally among GFP-trap resin (Chromotek) and fresh control resin for co-immunoprecipitation (IP) experiments. An equal volume of dilution buffer (20 mM HEPES [pH 7.5], 150mM NaCl, 0.5 mM EDTA, 1 cOmplete Mini EDTA-free Protease Inhibitor Cocktail tablet per 20 mL) to lysate was added to each sample then IPs were incubated 1 h at 4°C followed by three washes with dilution buffer. IPs were eluted by adding SDS loading buffer to the washed resin pellet and boiling for 5 minutes at 95°C.

### siRNA knockdown and DNA transfection

Cells were seeded on 6 well plates and used in knockdown assays the following day. AP-4 knockdown was achieved using ON-TARGETplus AP4E1 siRNA (J-021474- 05; Dharmacon) and tepsin knockdown was achieved using ON-TARGETplus C17orf56 siRNA (J-015821-17; Dharmacon). Control cells were treated with ON-TARGETplus non-targeting siRNA (D-001810-01; Dharmacon). Transfections of siRNA were carried out with Oligofectamine (Thermo Fisher Scientific) with a final siRNA concentration of

7.5 nM (for each AP-4, tepsin, and control siRNA treatment) in complete culture media and assayed 48 hours after transfection. Cells were re-seeded 24 hours after siRNA treatment to a lower cell density on 6 well plates (with or without coverslips) for Western blotting and immunofluorescence assays. For rescue experiments, cells were subjected to an additional transfection step. Transfection with siRNA was conducted as described above, but cells were seeded into and maintained throughout the experiment in antibiotic-free media (MEM-alpha with 10% FBS, no G418). 4 hours after incubation with siRNA treatment, wild-type and mutant tepsin-myc constructs were transfected using Fugene 4K (Promega) at 1.5:1 Fugene:DNA ratio following manufacturer protocol. Replating and a total time course of 48 hours from initial siRNA knockdown were maintained as described above.

### Fluorescence microscopy

Cells were seeded onto 12 mm #1.5 glass coverslips (Fisher Scientific) coated with Histogrip (Invitrogen). Coverslips were imaged with a Nikon Spinning Disk confocal microscope equipped with a Photometrics Prime 95B sCMOS monochrome camera; Plan Apo Lambda Oil 60x 1.40 NA WD 0.13mm objective; 405, 488, 561, and 647 nm excitation lasers. Image analysis was conducted using Nikon’s NIS Elements AR Analysis software GA3 pipelines.

For autophagic flux assays: mRFP-GFP-LC3B HeLa cells were fixed in 3% paraformaldehyde (Electron Microscopy Sciences) in PBS-CM (1X PBS with 0.1 mM CaCl_2_, 1 mM MgCl_2_) at room temperature for 20 minutes. Residual paraformaldehyde was quenched by incubating with 50 mM NH_4_Cl for 10 minutes. Coverslips were washed three times in PBS-CM with a final wash in Milli-Q H_2_O (Millipore Sigma). To preserve fluorophore signal, coverslips were kept in a dark box during all incubations. Coverslips were mounted in Prolong Diamond with DAPI (Invitrogen). Quantification of autophagic flux was performed using spinning disk confocal z-stack images. Individual cell regions of interest (ROIs) were generated using the “grow objects” function on the DAPI [excitation: 405 nm] signal. Punctate structures corresponding to autophagosomes (defined as having mRFP [excitation: 561 nm] and GFP [excitation: 488 nm] signal) or autolysosomes (mRFP signal only) were segmented using the “threshold” function then subjected to 3D analysis functions to obtain object counts and measure object volume. Data from each replicate was transformed using min-max normalization and individual data points from all replicates were combined for statistical analysis and data visualization.

For ATG9A/TGN46 immunofluorescence assays: mRFP-GFP-LC3 HeLa cells were either fixed and permeabilized in ice-cold methanol for 10 minutes at -20°C (tepsin-myc rescue experiments) or fixed in 3% paraformaldehyde PBS-CM followed by 5 min permeabilization with ice cold methanol at -20°C. Coverslips were washed three times with PBS-CM then blocked for 1 hour in 1% bovine serum albumin (BSA) in PBS-CM. Primary antibody incubations were conducted overnight at 4°C. The next day, coverslips were washed 3 times (10 minutes each) with BSA blocking buffer.

Fluorescent secondaries were diluted in BSA blocking buffer and incubated for 2 hours at room temperature protected from light. Coverslips were then washed two times with BSA block, once with PBS-CM, and once with dH_2_O before being mounted in Prolong Diamond with DAPI. Fixation effectively quenched exogenous mRFP-GFP-LC3B signal confirmed by comparing secondary antibody-only control cells to immuno-stained cells (Figure S3C). Quantification of ATG9A object measurements and colocalization with TGN46 was performed using spinning disk confocal z-stack images. Individual cell ROIs were generated using the 3D segmentation “OR” function to combine two ROIs (ROI-A and ROI-B) resulting in more precise cell segmentation. ROI-A was defined by the DAPI [excitation: 405 nm] signal subjected to the “grow objects” function and ROI-B was defined by the ATG9A channel [excitation: 647 nm] subjected to the “fast smooth” function (sigma=25 or sigma=18 [rescue experiments]). Combined ROIs were subjected to the “separate objects” function to generate final individual cell ROIs. ATG9A and TGN46 channels were background subtracted using the “detect regional maxima” function. The TGN was segmented using an intensity threshold on the TGN46 channel [excitation: 488 nm]. ATG9A objects within the TGN segmented area were defined as “*trans*-Golgi ATG9A” and ATG9A objects outside the TGN area were defined as “non-Golgi ATG9A”. The two populations of ATG9A objects were subjected to 3D measurement functions “max object intensity” and “object volume”. For Mander’s colocalization analysis, max intensity projections were generated for the ATG9A and TGN46 channels followed by background subtraction by the “detect regional maxima” function. For tepsin-myc rescue experiments, the mean intensity of the myc channel in each cell area was measured and used to subset the dataset in RStudio to specifically analyze transfected cells. Data across all replicates was combined for statistical analysis and data visualization.

For ATG9A/LC3 immunofluorescence assays: HeLa cells were fixed in 3% paraformaldehyde PBS-CM followed by 10 min permeabilization with ice cold methanol at -20°C. Quantification of ATG9A and LC3 colocalization was performed as described for ATG9A/TGN46 above with the following changes: All z-stack images were denoised using the denoise.ai feature in Nikon Elements software. Additionally, ROI-B for generating individual cell ROIs was defined by the LC3 channel [excitation: 488 nm] subjected to the Gaussian filter (sigma=10). For analysis of LC3 structures, LC3 objects were segmented using the “threshold” function then subjected to 3D analysis functions to measure object volume and object max fluorescence intensity. Data across all replicates was combined for statistical analysis and data visualization.

All statistical significance was analyzed using R with scripts written in RStudio and packages described above. Normalcy was assessed based on box-and-whisker plots for all data. These data did not exhibit a normal Gaussian distribution, so non-parametric tests were used. Nonparametric analysis of variance was assessed by the Kruskal-Wallis test and significant results were followed by a Dunn test with Bonferroni correction. When appropriate, two independent samples were compared using the Mann-Whitney U test. Data was graphed using ggplot2 functions in RStudio.

## Acknowledgements

We sincerely thank Margaret Robinson (Cambridge Institute for Medical Research) for providing the tepsin-GFP HeLa cell line and David Rubinsztein (Cambridge Institute for Medical Research) for providing the mRFP-GFP-LC3B HeLa cell line. We also thank members of the Jackson lab for helpful discussion and critical reading of the manuscript. NSW, JEG, CIC, and LPJ are supported by NIH R35GM119525. LPJ is a Pew Scholar in the Biomedical Sciences, supported by the Pew Charitable Trusts. NSW and CIC were partly supported by the Molecular Biophysics Training Grant NIH 5T32GM008320. JEG was partly supported by the Postdoctoral Program in Functional Neurogenomics NIH 5T32MH065215. Imaging and image analysis was performed in part using the Vanderbilt Cell Imaging Shared Resource (supported by NIH grants CA68485, DK20593, DK58404, DK59637 and EY08126).

## Author Contributions

Natalie S. Wallace: Investigation; Formal Analysis; Validation; Visualization; Writing- Original Draft; Writing- Review & Editing

John E. Gadbery: Investigation

Cameron I. Cohen: Investigation

Amy K. Kendall: Investigation

Lauren P. Jackson: Conceptualization; Writing- Original Draft; Writing- Review & Editing; Supervision; Project administration; Funding acquisition

---

**Figure S1:**
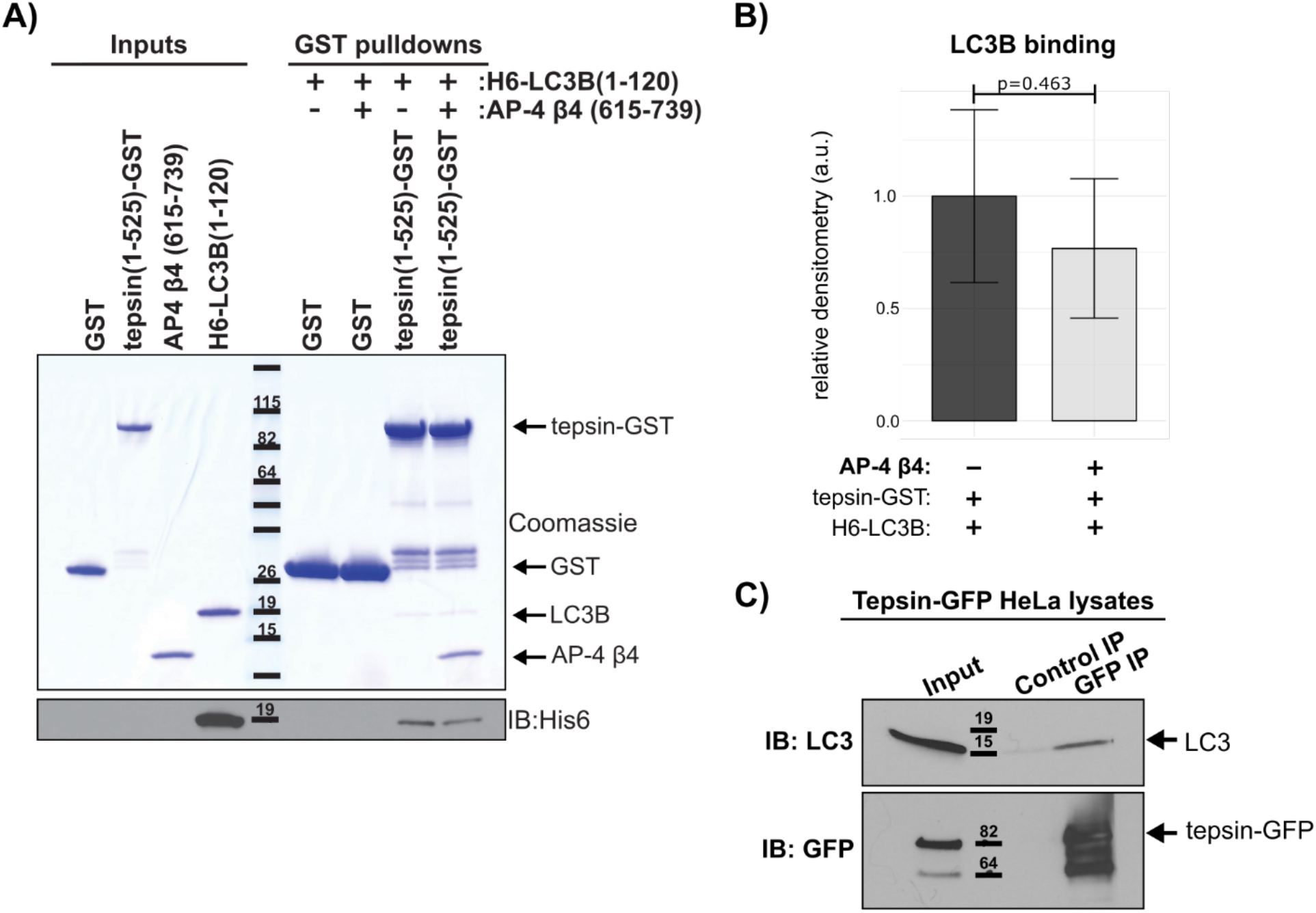
**Tepsin binds LC3B independently of AP-4-binding *in vitro.*** (A) Recombinant full-length tepsin(residues 1-525)-GST pulls down recombinant LC3B (residues 1-120)-His6x in the presence or absence of AP-4 β4 appendage domain. Binding between tepsin and LC3B was detectable on a Coomassie-stained SDS-PAGE gel and further confirmed by Western blot (α-His; Abcam ab184607). (B) Presence of AP-4 β4 appendage domain did not significantly change tepsin/LC3B binding *in vitro*. Bar plots (average ± standard deviation) depict quantification of LC3B binding to tepsin- GST in the presence or absence of AP-4 β4 appendage domain. Statistical analysis by Student’s t-test of three independent experiments with p-value indicated (p=0.463). (C) LC3 co-immunoprecipitates (IP) with tepsin from HeLa cells stably expressing tepsin-GFP detected by Western blot (α-LC3B: Abcam ab48394; α-GFP: Abcam ab6663). Control IP used unconjugated agarose resin. A and C: representatives of three independent experiments.

**Figure S2:**
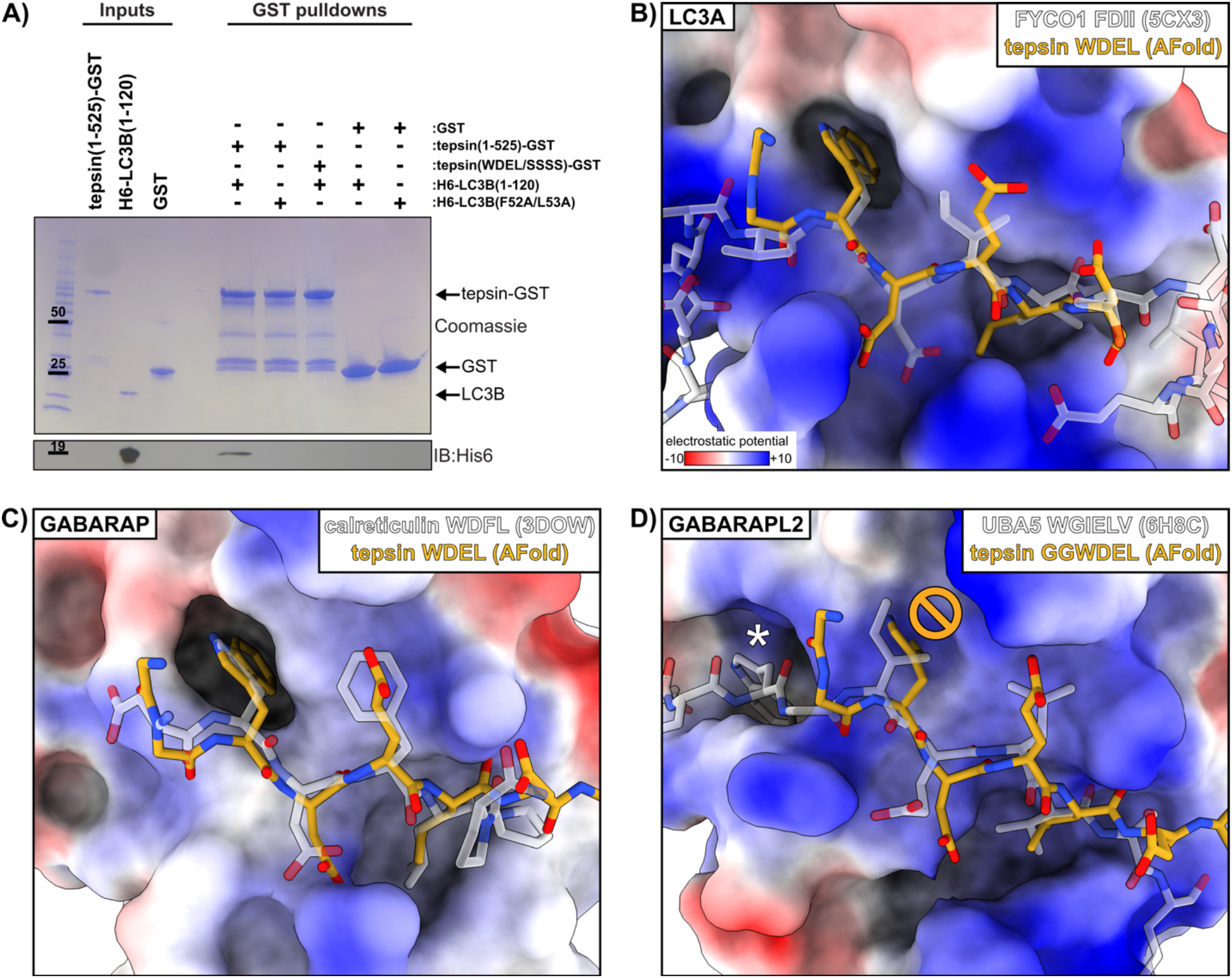
Biochemical experiments and computational modelling of the tepsin LIR motif explains binding selectivity among ATG8 family members. (A) Coomassie-stained SDS-PAGE gel and Western blot (α-His; Abcam ab184607) of GST pulldowns. Full-length wild-type tepsin (residues 1-525; WT) does not bind LDS mutant LC3B (F52A/L53A) and LIR mutant (WDEL/SSSS) tepsin does not bind wild-type or LDS mutant LC3B indicating the LIR motif substantially contributes to the tepsin/LC3B interaction *in vitro*. Free GST was used as a negative control. (B-D) Alphafold Multimer models were generated for mATG8 proteins and superposed with experimental structures: (B) LC3A, (C) GABARAP, (D) GABARAPL2. The hydrophobic residues of the LIR motif dock into hydrophobic pockets of LC3A and GABARAP, although the surface of GABARAP has reduced electrostatic potential compared to LC3; the tepsin Trp residue clashes with the surface in the experimental GABARAPL2 model (indicated) which has an elongated motif to dock the Trp residue in an alternative pocket (*).

**Figure S3:**
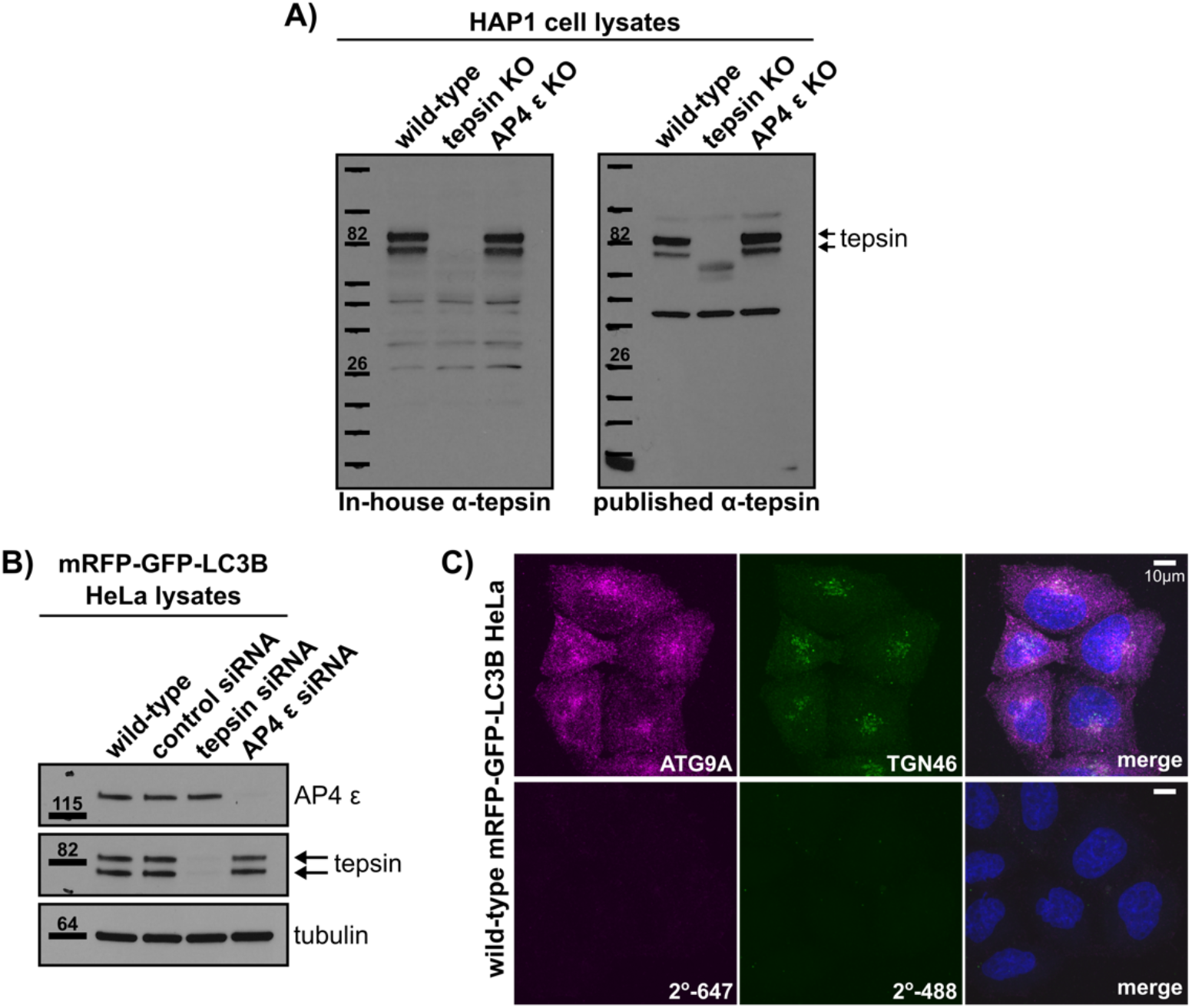
Antibody validation in HAP1 AP-4 ε or tepsin knockout cells and staining controls in mRFP-GFP-LC3B HeLa cells. (A) Manufactured tepsin antibody (reported here; described in methods) compared to a published tepsin antibody provided by the Robinson Lab in HAP1 knockout (KO) cells (Borner et al., 2012). Both antibodies predominantly recognize a doublet around 82 kDa which is absent in HAP1 tepsin KO cells. (B) Tepsin or AP-4 ε (gene: AP4E1) siRNA-treated mRFP-GFP-LC3B HeLa cells assayed by Western blot exhibit a corresponding loss of tepsin (α-tepsin; In-house see methods) or AP-4 ε (α-AP-4 epsilon; BD Transduction Labs 612019) respectively. Tubulin is presented as a loading control (α-alpha-tubulin; Proteintech 66031). (C) Exogenous GFP signal from mRFP-GFP-LC3B and TGN46 immunostaining are both visualized by a 520 nm emission peak (see methods). Representative maximum intensity projection confocal images taken from mRFP-GFP-LC3B HeLa cell line demonstrate immunostaining (see methods) effectively quenches exogenous GFP signal from mRFP-GFP-LC3B. Top row: immunostained for ATG9A (α-ATG9A Abcam ab108338; secondary α-Rabbit-647 Thermo Fisher Scientific A32733) and TGN46 (α- TGN46 Bio-Rad AHP500GT; secondary α-Sheep-488 Thermo Fisher Scientific A11015). Bottom row: incubated only with secondary fluorescent antibodies (α-Rabbit- 647 Thermo Fisher Scientific A32733; α-Sheep-488 Thermo Fisher Scientific A11015). The GFP signal from exogenous mRFP-GFP-LC3 does not contribute to TGN46 signal, validating analysis of TGN46-positive staining as *trans*-Golgi compartments in these cells. Scale bar: 10 µm.

**Figure S4:**
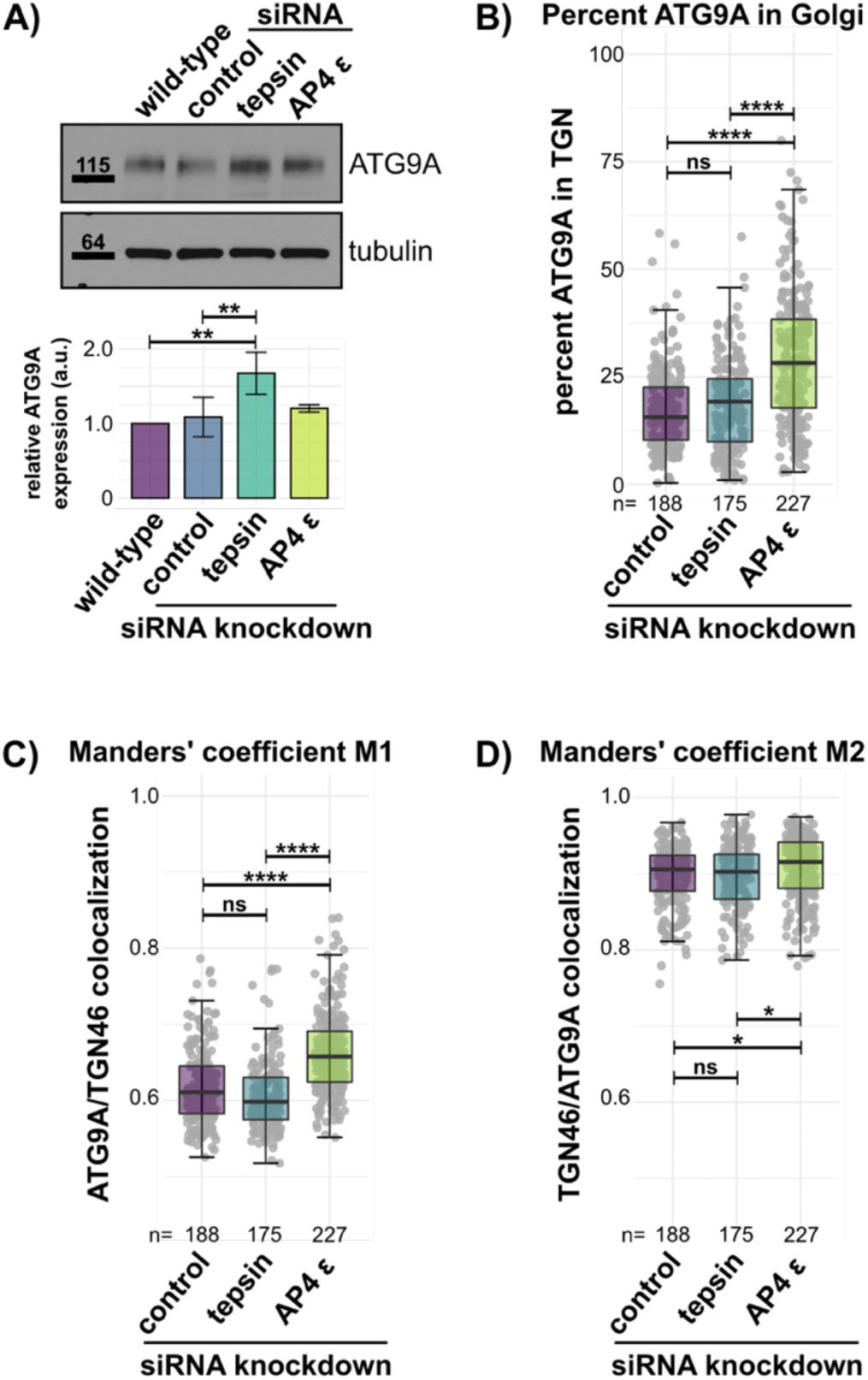
Tepsin depletion elevates ATG9A expression in mRFP-GFP-LC3B HeLa cells. (A) Representative Western blot for ATG9A (α-ATG9A Abcam ab108338) expression in tepsin- or AP-4-depleted lysates (knockdown shown in Figure S3). ATG9A expression is 1.5 times increased following tepsin-depletion. Bar plots (average ± SD) depict quantification of the ATG9A expression level relative to wild-type ATG9A expression. One-way ANOVA with Tukey post-hoc test; n=3. Tubulin was used as a loading control (α-alpha-tubulin; Proteintech 66031). (B) Percent of ATG9A signal in the TGN (defined by TGN46 signal) shows accumulation of ATG9A in the TGN within AP-4 ε-depleted cells. (C) Manders’ coefficient M1 indicating the fraction of ATG9A signal coincident with *trans*-Golgi marker, TGN46. (D) Manders’ coefficient M2 indicating the fraction of TGN46 signal coincident with ATG9A. These M1 and M2 results indicate the majority of ATG9A signal is retained within the TGN of AP-4 ε-depleted cells. The absolute ratio of ATG9A signal within the TGN is not significantly affected in tepsin- depleted cells. Quantification of four independent experiments (n=total cell count). Statistical results from Kruskal-Wallis test, Dunn test with Bonferroni correction; ns>0.05, *p≤0.05, **p≤0.01, ***p≤0.001, ****p≤0.0001.

**Figure S5:**
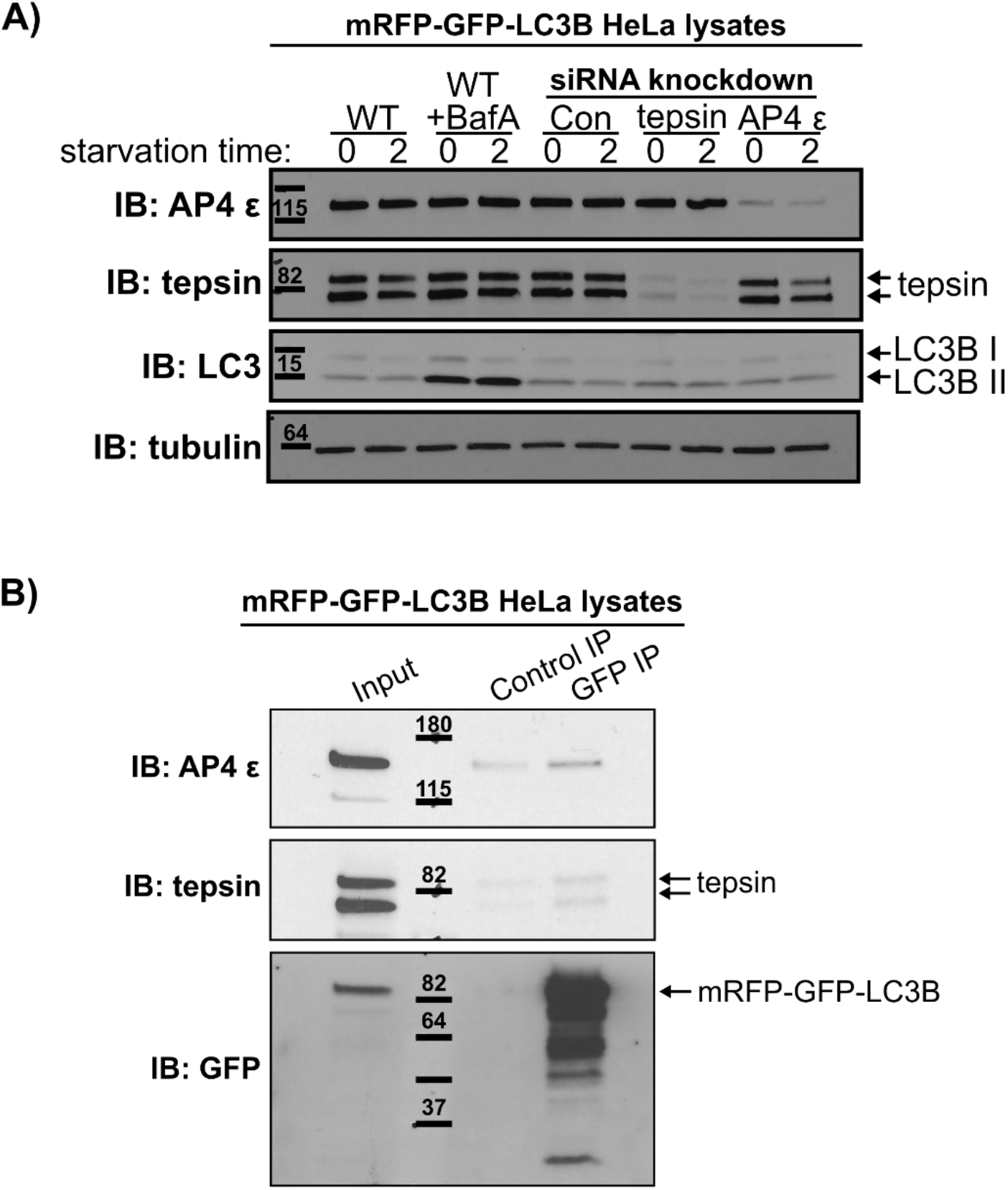
Tepsin and AP-4 interact with LC3B in mRFP-GFP-LC3B HeLa cells. (A) Representative Western blots from mRFP-GFP-LC3B HeLa cells treated with tepsin- or AP-4-targeting siRNA indicated effective tepsin or AP-4 knockdown. Western blots show LC3B accumulation following Bafilomycin treatment in wild-type cells confirming starvation conditions induced autophagy. Bafilomycin A treatment at 100 nM in indicated samples. (B) Western blots from co-immunoprecipitation experiments in mRFP-GFP-LC3B HeLa cell lysates. Endogenous tepsin co-immunoprecipitates on GFP-resin but not when using unconjugated resin (control). AP-4 ε also co-immunoprecipitates with mRFP-GFP-LC3B. Representative of 3 replicates. Antibodies: α-AP-4 epsilon, BD Transduction Labs 612019; α-tepsin, In-house (see methods); α- LC3B, Abcam ab48394; α-GFP, Abcam ab6663; α-alpha-tubulin, Proteintech 66031.

**Figure S6:**
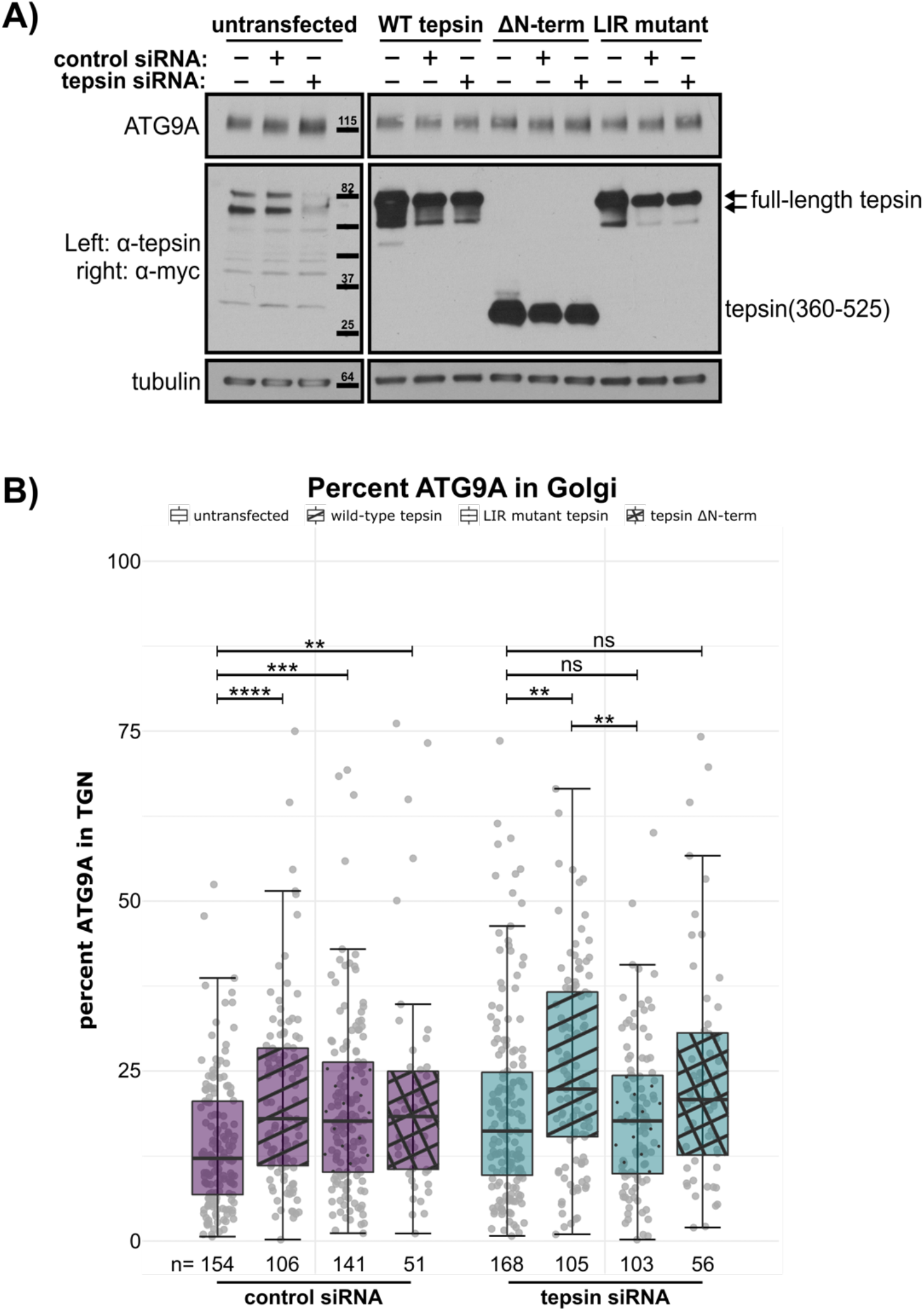
Expression of tepsin LIR mutant and ΔN-terminus tepsin constructs in tepsin depleted cells. (A) Representative Western blots from mRFP-GFP-LC3B HeLa cells treated with control or tepsin siRNA followed by transfection of myc-tagged tepsin constructs: wild-type tepsin, tepsin Δ N-terminus (residues 360-525; ΔΝ-term), or tepsin LIR mutant (WDEL/SSSS) as indicated. Western blots indicate siRNA-resistant transfection of each tepsin construct. Antibodies: α-ATG9A, Abcam ab108338; α-tepsin, In-house (see methods); α-myc-tag, Cell Signaling Technology 2276; α-alpha-tubulin, Proteintech 66031. (B) Percent ATG9A signal in the TGN (defined by TGN46 signal) indicates exogenous expression of tepsin constructs uniformly affects ATG9A cellular distribution in control-treated cells. Reintroducing LIR mutant or ΔN-term tepsin constructs following tepsin-depletion does not significantly affect ATG9A cellular distribution. Statistical results from Kruskal-Wallis test, Dunn test with Bonferroni correction; ns>0.05, *p≤0.05, **p≤0.01, ***p≤0.001, ****p≤0.0001.

**Figure S7:**
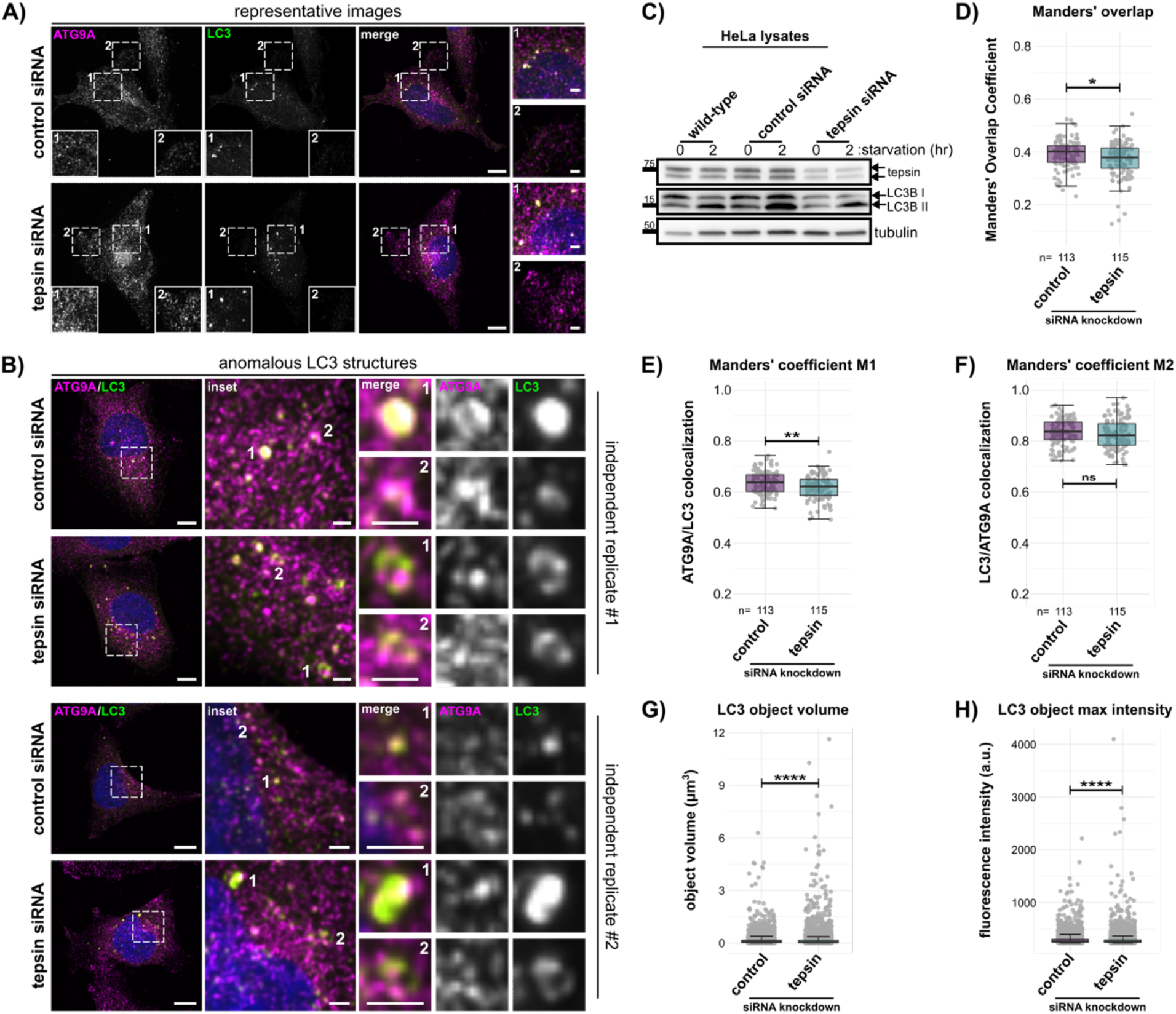
Tepsin-depleted HeLa cells accumulate ATG9A separately from LC3B structures. (A-B) Maximum intensity projection confocal images of basal HeLa cells immunostained for ATG9A (α-ATG9A Abcam ab108338; secondary α-Rabbit-647 Thermo Fisher Scientific A32733) and LC3 (α-LC3 MBL M152-3; secondary α-Mouse- 488 Thermo Fisher Scientific A32723). Cells were treated with non-targeting siRNA (control) or tepsin siRNA. Scale bar: 10 µm; inset scale bar: 2 µm. Representative images (A) show peripheral ATG9A accumulations do not co-stain for LC3 in tepsin- depleted HeLa cells. (B) A few tepsin-depleted cells, across independent replicates, exhibit anomalous LC3 structures not observed in control cells. (C) Representative Western blots from HeLa cells treated with control or tepsin siRNA. LC3BI and LC3BII levels were used to monitor autophagy levels in basal HeLa cells. Antibodies: α-tepsin, In-house (see methods); α-LC3B, Abcam ab48394; α-GFP, Abcam ab6663; α-alpha-tubulin, Proteintech 66031. (D) Manders’ overlap coefficient indicates on a whole cell level, less ATG9A co-localizes with LC3 after tepsin depletion. (E-F) The absolute ratios of ATG9A and LC3 coincidence indicates tepsin-depleted cells have a smaller fraction of ATG9A signal coincident with LC3 (E) while LC3 signal is similarly coincident with ATG9A (F) compared to control cells. Tepsin-depleted cells exhibit larger (G) and brighter (H) LC3 structures compared to control cells. Statistical results from Mann- Whitney U-test; ns>0.05, *p≤0.05, **p≤0.01, ***p≤0.001, ****p≤0.0001.

**Figure S8:**
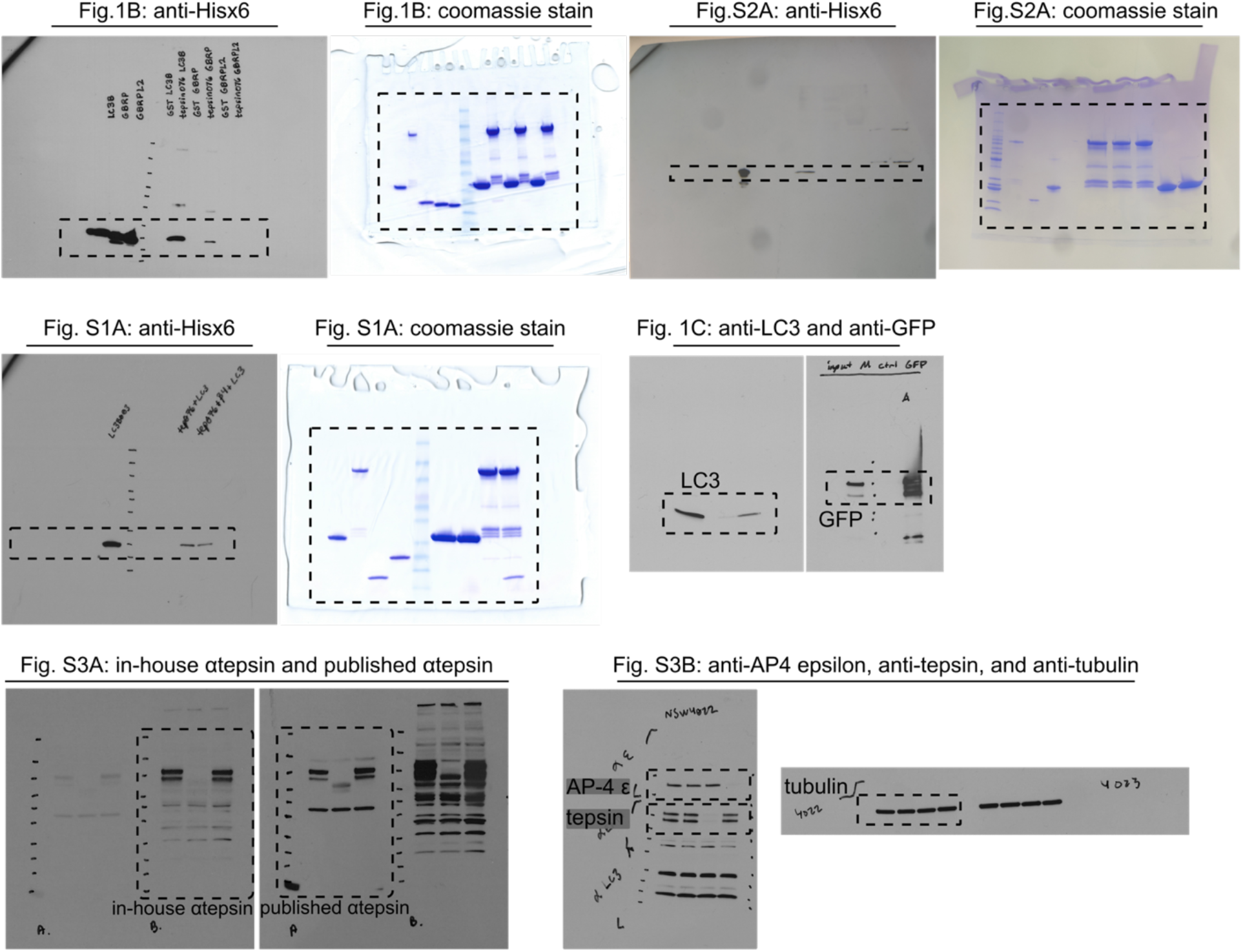
Uncropped Western blot films and Coomassie-stained SDS-PAGE gels from. Figure 1 **and Supplementary Figures S1, S2, and S3.** Dashed boxes denote regions cropped for the indicated figure.

**Figure S9:**
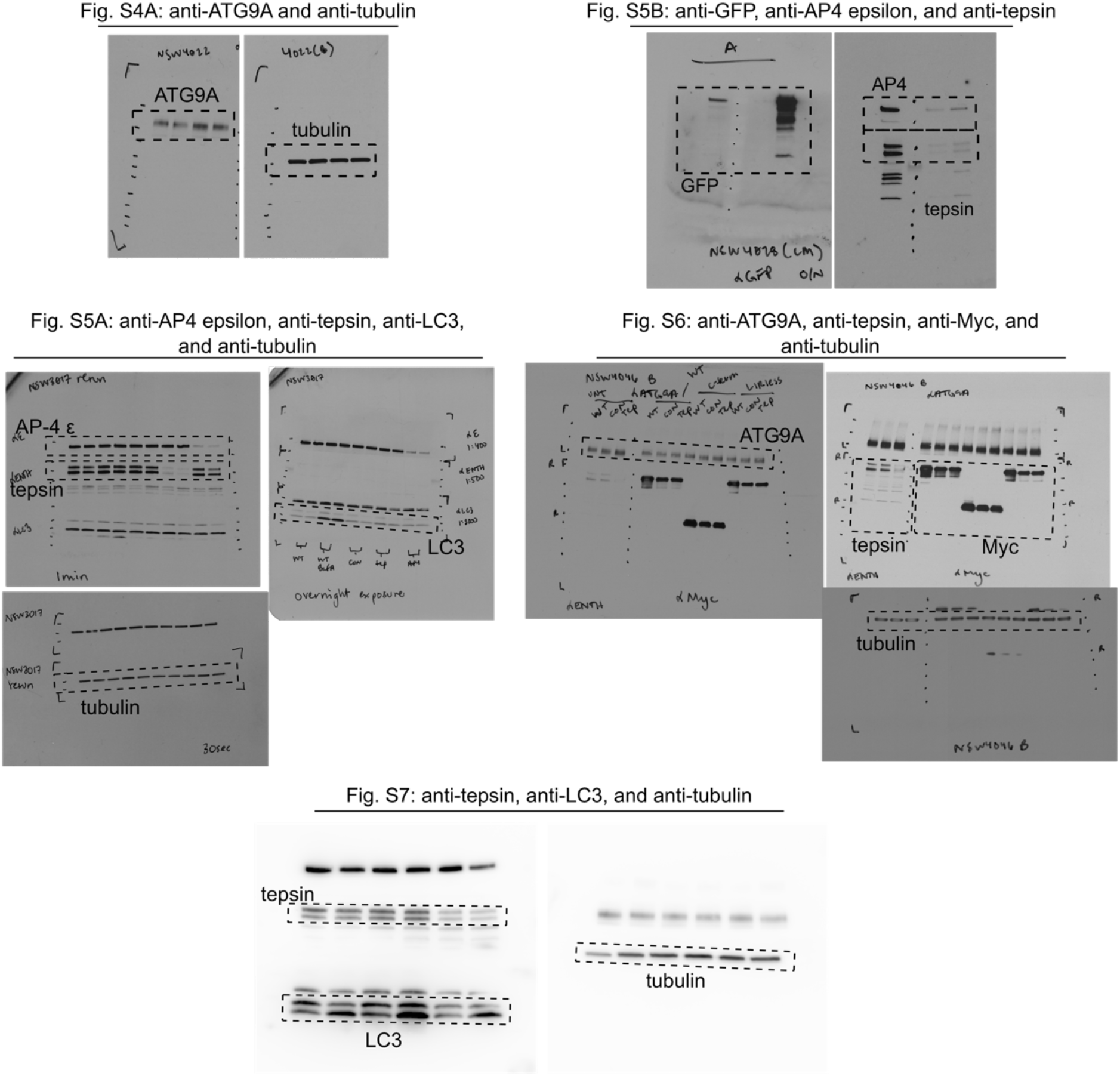
Uncropped Western blot films and Coomassie-stained SDS-PAGE gels from Supplementary Figures S4-S7. Dashed boxes denote regions cropped for the indicated figure.

